# Distinct transcriptomic cell types and neural circuits of the subiculum and prosubiculum along the dorsal-ventral axis

**DOI:** 10.1101/2019.12.14.876516

**Authors:** Song-Lin Ding, Zizhen Yao, Karla E. Hirokawa, Thuc Nghi Nguyen, Lucas T. Graybuck, Olivia Fong, Phillip Bohn, Kiet Ngo, Kimberly A. Smith, Christof Koch, John W. Phillips, Ed S. Lein, Julie A. Harris, Bosiljka Tasic, Hongkui Zeng

## Abstract

Subicular region plays important roles in spatial processing and many cognitive functions and these were mainly attributed to subiculum (Sub) rather than prosubiculum (PS). Using single-cell RNA-sequencing (scRNA-seq) technique we have identified up to 27 distinct transcriptomic clusters/cell types, which were registered to anatomical sub-domains in Sub and PS. Based on reliable molecular markers derived from transcriptomic clustering and in situ hybridization data, the precise boundaries of Sub and PS have been consistently defined along the dorsoventral (DV) axis. Using these borders to evaluate Cre-line specificity and tracer injections, we have found bona fide Sub projections topographically to structures important for spatial processing and navigation. In contrast, PS along DV axis sends its outputs to widespread brain regions crucial for motivation, emotion, reward, stress, anxiety and fear. Brain-wide cell-type specific projections of Sub and PS have also been revealed using specific Cre-lines. These results reveal two molecularly and anatomically distinct circuits centered in Sub and PS, respectively, providing a consistent explanation to historical data and a clearer foundation for future functional studies.

**Highlights:**

1. 27 transcriptomic cell types identified in and spatially registered to “subicular” regions.
2. Anatomic borders of “subicular” regions reliably determined along dorsal-ventral axis.
3. Distinct cell types and circuits of full-length subiculum (Sub) and prosubiculum (PS).
4. Brain-wide cell-type specific projections of Sub and PS revealed with specific Cre-lines.

**In Brief:** Ding et al. show that mouse subiculum and prosubiculum are two distinct regions with differential transcriptomic cell types, subtypes, neural circuits and functional correlation. The former has obvious topographic projections to its main targets while the latter exhibits widespread projections to many subcortical regions associated with reward, emotion, stress and motivation.

## Introduction

The subicular complex of the hippocampal formation has been reported to play important roles in many brain functions such as learning and memory, spatial navigation, emotion, reward, stress, motivation, and endocrine regulation (Aggleton and Christiansen, 2015; Herman and Mueller, 2006; O’Mara et al., 2009). The subicular complex is also heavily involved in many neurological and psychiatric diseases such as Alzheimer’s disease, temporal lobe epilepsy, schizophrenia, autism, anxiety disorder and drug addiction (Coras et al, 2014; Godsil et al., 2013; Van Hoesen and Hyman, 1990). To explore the anatomical substrates of these functions and diseases, neuroscientists have started to pinpoint specific subicular subfields for their cell types, neural circuits, physiological properties, and the effects of lesion or stimulation (Bienkowski et al., 2018; Cembrowski et al., 2018a; Huang et al, 2017; Preston-Ferrer et al, 2016; Tang et al., 2016). An important first step to characterize the subicular subfields is to accurately identify and target these subfields and their cell types.

The subfields of the subicular complex mainly include the prosubiculum (PS), subiculum proper (Sub or S), presubiculum [PrS, including postsubiculum (PoS), i.e. dorsal PrS (PrSd)] and parasubiculum (PaS) (Ding 2013; Rosene and van Hoesen, 1987). The concept of PS was proposed and refined by many neuroscientists (see Lorente de No, 1934; Rosene and Van Hoesen, 1987; Saunders et al., 1988a, b). Many previous studies in monkey have adopted the definition of PS, which is a narrow and oblique region between CA1 and Sub with strong AChE staining (Arikuni et al., 1994; Barbas and Blatt, 1995; Fudge et al., 2012; Saunders et al., 1988b; Wang and Barbas, 2018; Yukie, 2000). However, the term PS has not been fully accepted yet, especially in rodent literature, in which PS was often treated as part of Sub with a dropout of the term PS (see Ding, 2013, for review). Accordingly, inconsistent and even opposite results often exist in PS and Sub studies between monkey and rodent and across different research groups. For example, in some retrograde tracing studies, neurons in PS rather than Sub were shown to project to bed nucleus of stria terminalis (BST) and amygdala when the term PS was used in rat (Christensen and Frederickson, 1998; Howell et al., 1991) and monkey (Fudge et al., 2012; Rosene and van Hoesen; 1987; Saunders et al, 1988; Wang and Barbas, 2018). In contrast, when the term PS was not used, neurons in Sub were reported to project to BST and amygdala in other retrograde tracing studies of the rat (Kishi et al., 2006; Ottersen, 1982; Shi and Cassell, 1999; Veening, 1978; Weller and Smith, 1982). Similar situation was observed for projections from PS and Sub to ventromedial prefrontal cortex [PFvm, including prelimbic (PL) and infralimbic (IL) cortices] and ventral striatum [VS, including nucleus accumbens (ACB) and olfactory tubercle (OT)]. For instance, after retrograde tracer injections into IL or VS, labeled neurons were mostly found in PS (Jay et al., 1989) or in “proximal Sub” (close to CA1; roughly corresponding to PS) rather than in “distal Sub” (close to PrSd) of the rat (Christie et al, 1987; Ishizuka, 2001; Phillipson and Griffiths, 1985;Witter, 2006; Witter et al., 1990). All above findings suggest the existence of PS as a distinct entity from Sub, with distinct outputs. If it is confirmed that PS rather than Sub projects to amygdala, VS, BST and IL, structures heavily involved in emotion, reward, stress and motivation (Aggleton and Christiansen, 2015; Herman and Mueller, 2006; O’Mara et al., 2009; Strange et al., 2014), then it is reasonable to hypothesize that it is PS rather than Sub that plays important roles in these brain functions and related diseases. Clarification of this issue will lead to clearer picture of region- and cell type-specific circuits of PS and Sub, to facilitate more accurate functional studies.

We have taken several approaches in this study to test and verify the hypothesis that PS exists as a distinct region and has distinct cell types and neural circuits in mice. First, we use unbiased hierarchical clustering of scRNA-seq transcriptomic profiles to identify cell types, showing that distinct molecular cell types exist in Sub and PS. Second, using region-specific gene markers we consistently delineate and update the boundaries of Sub and PS along dorsal-ventral (DV) axis. Third, with these boundaries as a guide we reveal differential afferent and efferent connections of Sub and PS at whole brain level. Fourth, utilizing different Cre-lines, we trace brain-wide projections of major cell classes in Sub and PS. Together, we have systematically linked distinct cell types, molecular signature, connectivity and anatomy of Sub and PS, and opened the potential to specifically target cell types and related circuits in future studies to understand related functions and diseases as mentioned above.

## Results

### Transcriptomic taxonomy of glutamatergic neurons in Sub and PS

Since recent molecular and connectional studies had not reached a full consensus regarding cell types and connectivity patterns of the mouse subicular regions (Bienkowski et al., 2018; Cembrowski et al., 2018a, 2018b), we first performed a transcriptomic survey of 17,062 glutamatergic cells isolated from the subicular complex (see Star*Methods). In addition, the rodent homolog of monkey and human hippocampo-amygdaloid transition area (HA; see Ding and Van Hoesen, 2015; Rosene and Van Hoesen, 1987) is also located in this region, mainly ventral to PS (see Allen Mouse Brain Common Coordinate Framework, CCFv3; Atlas.brain-map.org). We performed two small micro-dissections (including regions PrS-PoS-PaS and Sub-PS-HA) and a larger one (including all of PrS-PoS-PaS-Sub-PS-HA) along DV axis of the hippocampal formation (see Star*Methods and Figure 1A). Cells from the first two micro-dissections were sequenced with the SMART-Seq v4 method (Tasic et al, 2018), which only allows small sample sizes and was mainly used for confirmation of the 10X Chromium clustering results in this study, while those from the larger micro-dissection were sequenced with the 10X Chromium v2 method, which allows much larger sample sizes.

**Figure 1.**
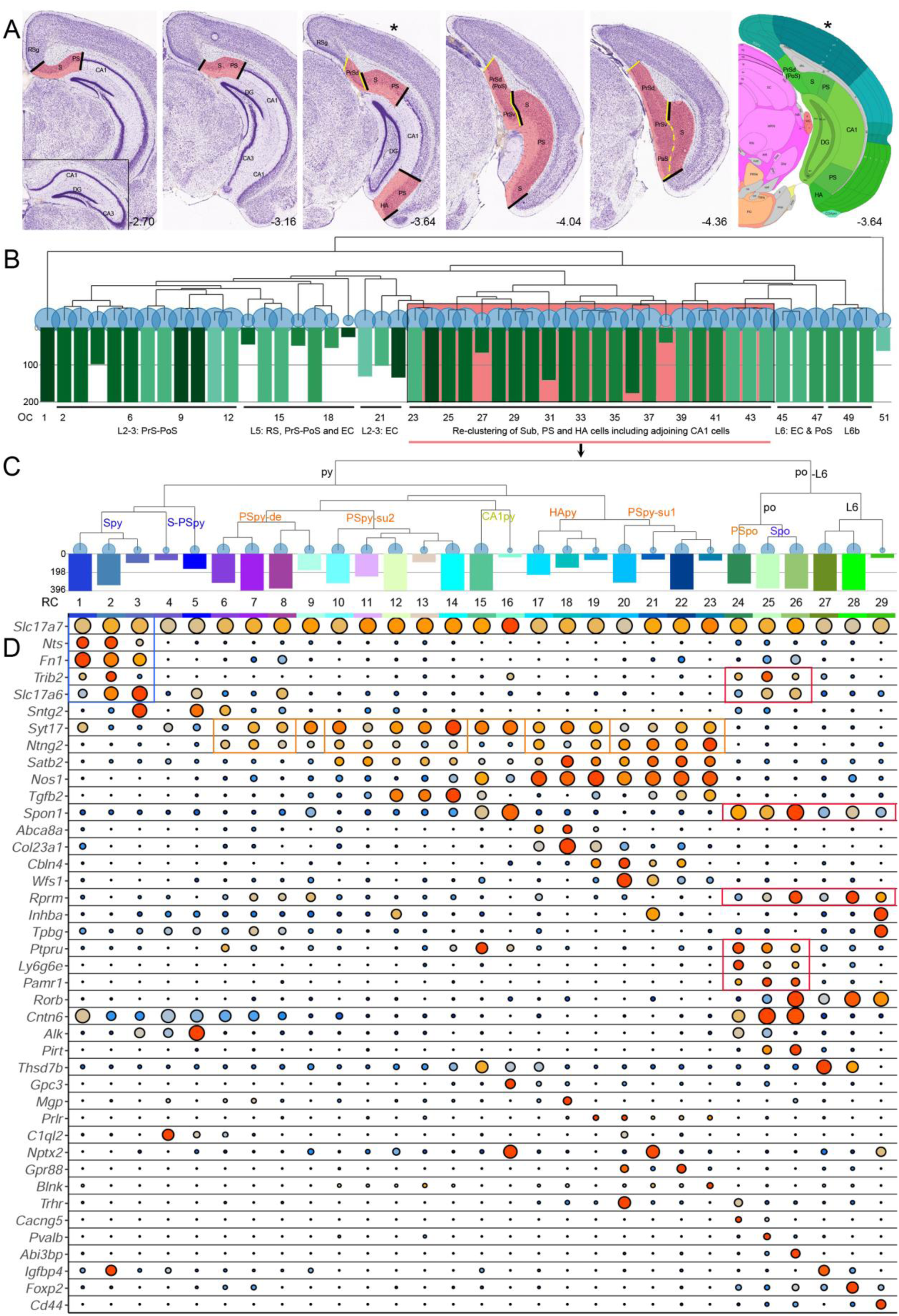
Transcriptomic classification of Sub and PS cell types. (**A)** Microdissection of tissue from Sub-PS-HA (region between black bars), PrS-PoS-PaS (region between yellow bars) or all subicular complex (all highlighted area) in sequential coronal sections. The inset in the first image shows lack of Sub-PS at more anterior levels. The last image shows the reference atlas plate for Sub-PS-HA region that matches the third image (*). Coordinates from Paxinos and Franklin (2012) are indicated at the bottom of each panel. (**B)** Unbiased hierarchical clustering of cells from subicular complex (10X). OC1-51 indicate corresponding original clusters (OC). Highlighted OC23-44 were re-clustered in C. (**C)** Hierarchical re-clustering of cells from Sub-PS-HA region (i.e. cells from OC23-44 in B). RC1-29 indicate corresponding new cluster numbers from re-clustering (RC). Note that PSpy is closer to CA1py and HApy than to Spy at transcriptional level. (**D)** Dot plot of selected gene markers (from 10X data) from re-clustering (see Table S1). Color of the dots indicates cluster mean expression (log scale), normalize by row by dividing the maximum value per row, so that the maximum normalized value is one. Blue to red corresponds to normalized values from zero to one. Size of the dots indicates % cells with CPM > 1. Major clusters and marker genes are outlined with colored boxes. S, subiculum; PS, prosubiculum; RSg, granular part of retrosplenial cortex; HA, hippocampo-amygdaloid transition area; PrSd and PrSv, dorsal and ventral presubiculum; PoS, postsubiculum; PaS, parasubiculum; L2-3, L5, L6 and L6b, layers 2-3, 5, 6 and 6b of related cortices; py, pyramidal layer; po, polymorphic layer; Spy, pyramidal layer of the subiculum; Spo, polymorphic layer of the subiculum; PSpy, pyramidal layer of the PS; PSpy-de, deep portion of the pyramidal layer of PS; PSpy-su1, the most superficial portion of the pyramidal layer of PS; PSpy-su2, superfitial portion of the pyramidal layer of PS; PSpo, polymorphic layer of PS; CA1py, pyramidal layer of hippocampal field CA1; HApy, pyramidal layer of the HA. For other abbreviations see Table S3.

We performed a consensus clustering to combine the SMART-Seq and 10X datasets. For this study, we calculated a dendrogram specifically for the glutamatergic class of neurons which include 2,182 SMART-Seq cells and 14,880 10X cells. For visualization, 10X cells were down-sampled to up to 200 cells per cluster (original cluster, OC; Figure 1B). From this dendrogram, based on marker gene localization (Figure S1A and Table S1) we identified three major branches which correspond respectively to principal cells in (1) layers 2-3 and 5 of PrS-PoS-PaS and adjoining retrosplenial (RS) and entorhinal cortex (EC) regions (OC2-22, but not OC23 and OC24), (2) pyramidal cell layer of Sub-PS-HA and adjoining CA1 (OC23-38) and (3) deepest layers of all related regions (OC39-50). OC1 (for dentate gyrus cells) and OC51 (for Cajal-Ritzius cells) are two outliers (due to imperfect tissue micro-dissections, cells from neighboring regions are often found in scRNA-seq datasets). OC2-22 of the first major branch (at the left of Figure 1B), together with OC1 and OC51, were excluded from further analysis because they do not contain cells from our focused Sub-PS-HA region in this study. OC23 and OC24 of the first branch were located in the most superficial pyramidal layer of PS (PSpy). The second major branch (OC25-38) was spatially registered to deep PSpy (OC25, OC26), pyramidal layer of Sub (Spy; OC27-31), superficial pyramidal layer of HA (HApy) and PSpy (OC32-35), as well as pyramidal layer of adjoining CA1 (CA1py; OC36-38). The third major branch (OC39-50) was spatially registered to polymorphic layer of Sub and PS (Spo and PSpo; OC39-41), layer 6 (L6) of all related regions (OC42-47), and layer 6b (L6b, sometimes called layer 7) of related regions (OC48-50). Layers 6 and 6b were named because they are continuous with layers 6 and 6b of adjoining cortices, respectively, and because they are separate from the polymorphic layer at transcriptomic and anatomic levels (e.g. Figs. 1B; 2S, T; S1G).

To explore whether more refined clusters could be revealed from the Sub-PS-HA region, we pooled all cells (n = 8648) from OC23-38 and OC39-44, and performed hierarchical re-clustering (re-cluster, RC; Figures 1C; S2A). This resulted in 25 clusters in the Sub-PS-HA region, 3 clusters in adjoining CA1 and one cluster in the deep L6 of PrS (Figures 1C-D; 2A-T; S1B-N). Note that L6b cells (OC48-50) and L6 cells from the medial and lateral EC (OC45 and OC46) and PrS-PoS (OC47, superficial L6) were not included in this re-clustering. Compared with the original clustering, re-clustering revealed a few more clusters/cell types in PSpy and HApy but not in Spy and adjoining CA1py (Table S1). Interestingly, in both clustering, PSpy is closer (more similar) to CA1py and HApy than to Spy by gene expression distance (Figure 1B, C). In addition to dendrogram, we also performed tSNE-based nonlinear dimensionality reduction for visualization of these 29 clusters from re-clustering (see Figure S2B). Generally, six major subclasses can be identified in the Sub-PS-HA region (Figure 1C and Table S1), corresponding to Spy (RC1-3), deep PSpy (RC6-8), superficial PSpy (RC10-14), HApy (RC17-19), most superficial PSpy (RC20-23), Spo and PSpo (RC24-26), and L6 of the Sub and HA (RC28-29). Cells from two small clusters (RC4, RC5) lie in the superficial pyramidal layer at the border between ventral Sub and PS. Therefore, taking away adjoining CA1 (RC9, RC15, RC16) and PrSd (i.e. PoS; RC27) and adding L6b of Sub and PS from original clustering (OC48, OC49), a total of 27 clusters or cell types were revealed in the Sub-PS-HA region. These cell types were partially covered in the overall “subiculum” region sampled in Cembrowski et al. (2018b). The latter study revealed 8 clusters (with an additional one in CA1) which mainly represent coarse subclasses probably due to a smaller sample size used (n = 1150 cells) (for comparison of the clusters, see Table S1 and Figure S2A).

**Figure 2.**
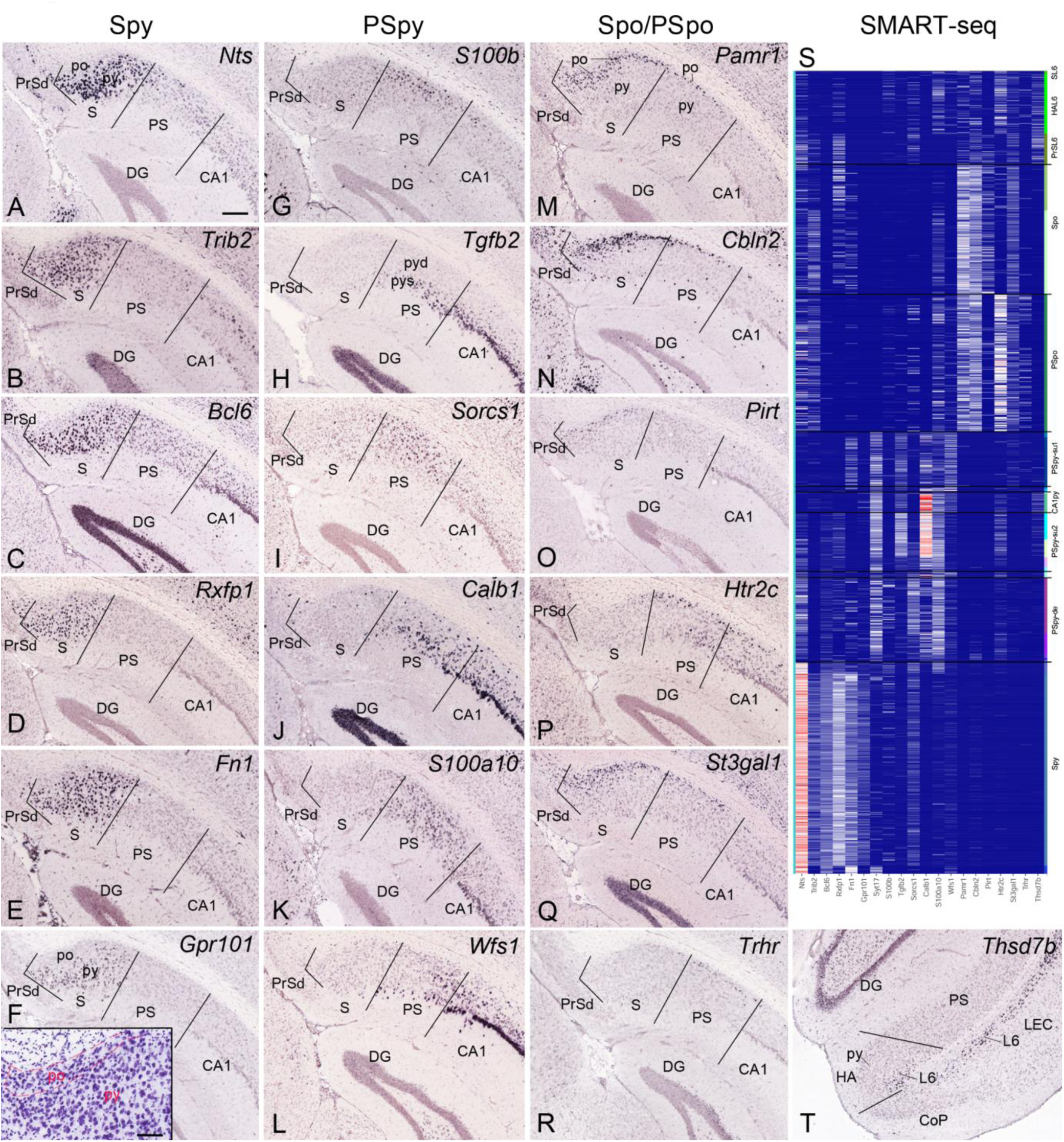
Major transcriptomic clusters mapped to anatomic regions. Black lines mark the areal borders in each panel. **(A-F)** Marker genes selectively or predominantly expressed in with no or few expressed cells in PSpy. The inset in (**F)** shows the different cell shapes and sizes between py and po. (**G-L)** Marker genes selectively or predominantly expressed in PSpy with no or few expressed cells in Spy. (**M-R)** Marker genes selectively or predominantly expressed in Spo and/or PSpo with no or few expressed cells in Spy and PSpy. (**S)** corresponding transcriptomic clusters revealed with SMART-seq, which is largely consistent with the results from 10X in Figure 1D. (**T)** Thsd7b expression in layer 6 (L6) of HA and EC. Bars: 210µm in A (for A-R and T); 95µm in the inset in F. For abbreviations see Table S3.

### Anatomical mapping of transcriptomic clusters

To facilitate accurate targeting and manipulation of specific cell types, we tried to correlate transcriptomic clusters with anatomical locations. For this we performed a survey of Allen Brain Atlas (ABA, Lein et al, 2007) *in situ* hybridization (ISH) dataset using distinct gene markers revealed at different branch levels (Figures 1D, S1B; Table S1). As shown in Figures 1-3, *Nts, Fn1, Rxfp1, Adcyap1, Gpr101* and *Bcl6* are expressed in most neurons located in anatomically defined Spy region, while genes such as *Ntng2* and *Syt17* are expressed in most neurons located in PSpy. Newly defined HA (Allen Mouse Brain CCFv3) also displays region-specific or region-enriched expression of genes such as *Col23a1, Rab3b, Id4, Abca8a, Gpc3, Unc5d, Lpl* and *Car10* (e.g. Figure S1C-F). HA was previously treated as ventral Sub or ventral CA1 in rodent (Bienkowski et al., 2018; Paxinos and Franklin, 2012) but has recently been found to be the homolog of monkey and human HA (Allen Mouse Brain CCFv3; Ding and van Hoesen, 2015; Rosene and Van Hoesen, 1987). HA is characterized by densely packed and modified pyramidal neurons in its superficial layer (HApy) and less densely packed small neurons in its deep layers (HAL6; Figure S1C-F). The polymorphic layers of Sub and PS (Spo and PSpo, respectively) show region-specific expression of *Ly6g6e, Cntn6, Pamr1, Cbln2* and *St3gal1* (Figures 1D; 2M, N, Q) while the deeper layer (layer 6) of Sub and HA expresses another set of region-specific genes such as *Sema3e, Thsd7b* and *Nppc* (Figure 3S, T; PS appears to lack this layer or has only scattered cells). Finally, a layer of cells lined at the gray-white matter border (layer 6b) of PrSd (i.e. PoS) and Sub were also revealed in this study (Figure S1G).

**Figure 3.**
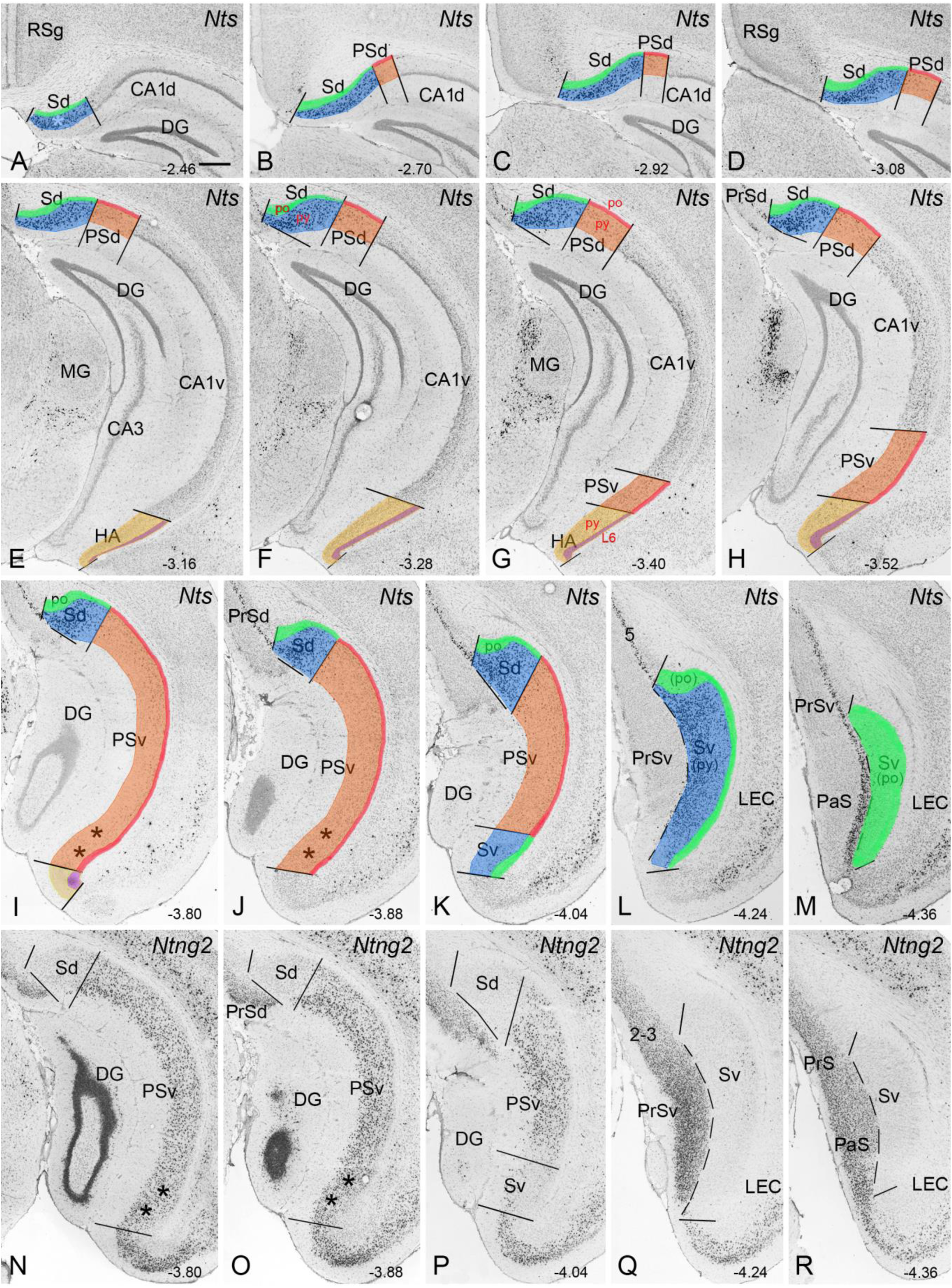
Borders, extent and topography of dorsal and ventral Sub and PS. (**A-M)** Borders, extent and topography of dorsal Sub and PS on *Nts* sequential rostral-caudal ISH sections. *Nts* is predominantly expressed in the pyramidal layer (py) of dorsal Sub (Sd in A-K) and ventral Sub (Sv in L) with no or few expressed cells in dorsal and ventral PS (PSd and PSv). Note that the polymorphic layer (po) of both dorsal and ventral Sub does not express *Nts*. (**N-R)** Borders, extent and topography of ventral Sub and PS on *Ntng2* sequential ISH sections. *Ntng2* is predominantly expressed in PS. Note that the ventral PS (PSv) is easily distinguishable from ventral Sub (Sv). The extent (width) of Sub decreases from dorsal (A) to ventral (M) while opposite is true for PS. The most dorsal part (* in A) contains only Sub while the most ventral part (** in I, J, N, O) contains only PS (this finding is obviously observed in sequential sagittal sections shown in Figure S4). Note also that the color-coded po of Sub and PS in Fig. 3A-M contains cells from deeper layers L6-L6b (for Sub) or L6b (for PS) for concise illustration. Bar: 350µm in A (for A-R). Coordinates from Paxinos and Franklin (2012) are indicated at the bottom of each panel. For abbreviations see the list in Table S3.

Three clusters (RC1-C3) could be further identified in the Spy. RC1 and RC2 correspond to distal Spy (away from PS) while RC3 to proximal Spy (close to PS). The gene markers for RC1 cells include *Cntn6, Dio3, Npsr1, Angpt1, St8sia2* and *Pdzrn4* while those for RC2 include *Igfbp4, Cyp26b1, Scn4b* and *Whrn* (e.g. Figures 1D; S1H, I; Table S1). Representative marker genes in RC3 include *Sntg2, S100b, Col6a1, Alk* and *Glra3* (Figures 1D; S1J). RC4 is a small cluster with cells located in the superficial pyramidal layer of the ventral Sub bordering PS and expresses genes *Teddm3, Luzp2, Galnt14, Fam19a1* and *C1ql2* (Figure S1K; Table S1). Another small cluster, RC5 has its cells in the superficial S-PSpy border region and expresses marker genes *Eps8, Dcc, Gpr101, Cyp26b1, Whrn, Alk, Gpc5* and *Glra3* (e.g. Figure S1L-N; Table S1).

In contrast to Spy, PSpy contains many subtypes (12 clusters), representing highly heterogenous cell types which often show laminar organization (Table S1; Figures 2; S3). RC6-8 and RC10-14 represent deep (PSpy-de) and superficial PSpy (PSpy-su2), respectively, while RC20-23 were mapped to the most superficial PSpy (PSpy-su1; Figures 2G-L; S3). *Cbln4* and *Nos1* are the representative gene markers for PSpy-su1 (RC20-23; see Figure S3B-F), which is generally located superficial to PSpy-su2 (RC10-14). The latter expresses another set of genes such as *Dlk1* and *Col25a1* (Figure S3G-K, S-U). This is consistent with the non-overlapping expression of *Cbln4* and *Dlk1* reported in a recent study (Cembrowski et al., 2018b). However, some genes (e.g. *Tgfb2* and *Satb2*) are expressed in both groups (e.g. Figure S3A, L-P). Moreover, gene expression difference along DV axis is detected in the PSpy-de group (RC6-8). For example, *S100b* and *Slc17a6* tend to be expressed strongly in dorsal portion (RC8; Figures 1D; 2G; 2S) while *Syt10* and *Mgp* tend to be expressed strongly in the ventral portion of PSpy-de (RC6-7; Figure S3A). In PSpy-su1 (RC20-23) and PSpy-su2 (RC10-14) subclasses, differential gene expression along DV axis is also found (see Table S1).

Genes selectively or predominantly expressed in PSpo (RC24; e.g. *Cacng5, Htr2c, Trhr* and *Gdpd2*) or Spo (RC25-26; e.g. *Plcb4, Pirt, Abi3bp* and *Pdzrn4*) were also observed, although more genes were expressed in both PSpo and Spo (Figures 1D, 2M-S). The latter genes include *Pamr1, Cbln2, Ly6g6e, Chrm2, Tle4, Kcnmb4, St3gal1, Trp53i11* and *Drd1a* (e.g. Figure 2M, N, Q). Finally, genes expressed in L6 (RC 28-29) and L6b (OC48-49) of Sub, PS and HA were often seen to extend to adjoining PrS (RC27, OC50) and EC. L6 cells express genes such as *Sema3e, Thsd7b, Car10* and *Nppc*, and HA has the thickest L6 which selectively or predominantly expresses genes *Sema3d, Foxp2, Thsd7b* and *Car10* (e.g. Figures 1D; 2T; S1F). L6b is just a single cell layer at the gray-white matter border expressing *Nxph4, Cplx3* and *Ctgf* (e.g. Figure S1G).

### Precise borders, topography and extent of Sub and PS

The borders of Sub, PS, HA and adjoining CA1 were not consistently defined previously in literature and commonly used brain atlases (e.g. Bienkowski et al., 2018; Cembrowski et al., 2018b; Paxinos and Franklin, 2012). Since we identified reliable and differential markers for these regions at transcriptional level, we next delineated precise boundaries between these regions along their DV axis with a combined use of the identified selective gene markers in sequential coronal (Figure 3) and sagittal (Figure S4) sections. For convenient description, Sub, PS and CA1 are roughly subdivided into dorsal (d) and ventral (v) parts since no clear markers are available for the DV borders. In coronal sections, the DV border between Sub and PS was placed at the dorsal edge of the most caudal PS (for Sd and Sv, see Figure 3K) or at the dorsal edge of the most caudal CA1 (for PSd and PSv, see Figure 3H). The locations of Sd, PSd, CA1d, Sv, PSv and CA1v as well as their topography were shown on sequential coronal sections stained for *Nts* (Figure 3A-M) and *Ntng2* (Figure 3N-R), which show complementary expression patterns. Sd medially adjoins granular part of the retrosplenial cortex (RSg) at dorsorostral levels (Figure 3A-E) and PrSd at ventrocaudal levels (Figure 3F-K). Laterally, Sd adjoins PSd at rostrodorsal levels (Figure 3A-H) and PSv at caudoventral levels (Figure 3I-K). Sv adjoins PrSv medially, PSv rostrally, LEC laterally and MEC caudally (Figure 3K-M). PSd is located lateroventral to Sd and mediodorsal to CA1d, while PSv is located rostral to Sv and ventral to CA1v. Both CA1v and PSv are connected with cortical amygdalar nucleus via HA (Figures 3E-H and S4A, G-I). From dorsorostral to ventrocaudal levels, the width and extent of Sub decreases while that of PS increases. Based on the present delineation it is surprising to find that the previously defined Sv in fact belongs to PSv (see the region marked with ** in Figure 3I, J, N, and O), because this region expresses many typical PS (e.g. PSd) genes including *Ntng2, Calb1, Nnat, Syt17* and *Adra1a* but not typical Sub (e.g. Sd) genes revealed in this and recent studies (Bienkowski et al., 2018; Cembrowski et al., 2018). The real Sv only appears at the most caudal levels of the coronal sections (Figure 3K-M, P-R). Finally, it is also worth mentioning that the borders between Sub and PS and between PS and CA1 are oblique with variable orientation at different levels (Figure 3A-M), making it very difficult to restrict neuronal tracer or drug injections in only one region. All these findings were confirmed in sequential sagittal sections stained for *Calb1* ISH (marker gene for PS, Figure S4A-H) and *Bcl6* ISH (marker gene for Sub, Figure S4I-O). Finally, it is interesting to find that PSv and Sv occupy the superficial and deep pyramidal layers, respectively, at level H (*Calb1*) or level P (*Fn1*, another marker for Sub, Figure S4P) of the sagittal sections.

### Distinct brain-wide projection patterns of the Sub and PS

To determine whether the Sub and PS have overall similar, distinct or mixed projection patterns, we compared brain-wide projection targets of Sub and PS by separating Sub- and PS-injected cases. Since most of previous connectional studies used wild-type animals and traditional neuronal tracers, we have mostly made use of Cre-driver mice (Table S2) in this study to avoid potential fibers of passage issue and to make evaluation of injections and interpretation of results easier. As expected, it is extremely difficult to restrict anterograde tracer injections (rAAV, see Methods) in Sub or PS of wild-type mice without leakage into adjoining PS or Sub. Among many cases with injections involving Sub of wild-type mice, we identified only 4 cases with injection sites mainly in the Sub (but not in PS) (Table S2). However, in Cre-driver mice, selectively targeting Sub is much easier. For example, in *Trib2*-F2A-CreERT2 and *Grik4*-Cre mice, where the gene driving Cre is expressed only in Sub but not PS (e.g. Figure 2B), the effective injection site would only be in Sub even if the injection covers both sub and adjoining PS because Cre-dependent GFP expression is only present in Cre-expressing neurons. In this study, we identified 16 Cre-mice with effective injection sites in Sub but not PS. The overall distribution of labeled axon terminals in the target regions of Sub injections is shown in Table S2. Briefly, Sd injections resulted in strong axon terminal labeling in RSg, PaS, PrS, Pro, MEC, MM, AV, AM and Re with weaker labeling in LS (e.g. Figures 4A, S5A1-I1; Table S2; for regional terminology and abbreviations see Table S3). In Sv-injected cases, labeled axon terminals were clearly found in the same target regions as in Sd cases (but at differential DV locations except in Re): RSg, PaS, PrS, Pro, MEC, AV, AM, MM (strong), Re and LS (weaker) (Figures 4B-D; 5A-O; S5A2-I2). In Re, projections from both Sd and Sv converge in the dorsolateral part (Figure S5J1 & J2). In all Sub-injected cases, no or few labeling was detected in PL-IL, PRC, VS (ACB+OT), BST, AON, AOB, amygdaloid nuclei and most of the hypothalamic regions excluding MM (Figures 4, 5, S5; Table S2). Taken together, the main targets of Sub include RSg, PaS, PrS, Pro, MEC, AV, AM, MM and Re (target set A).

**Figure 4.**
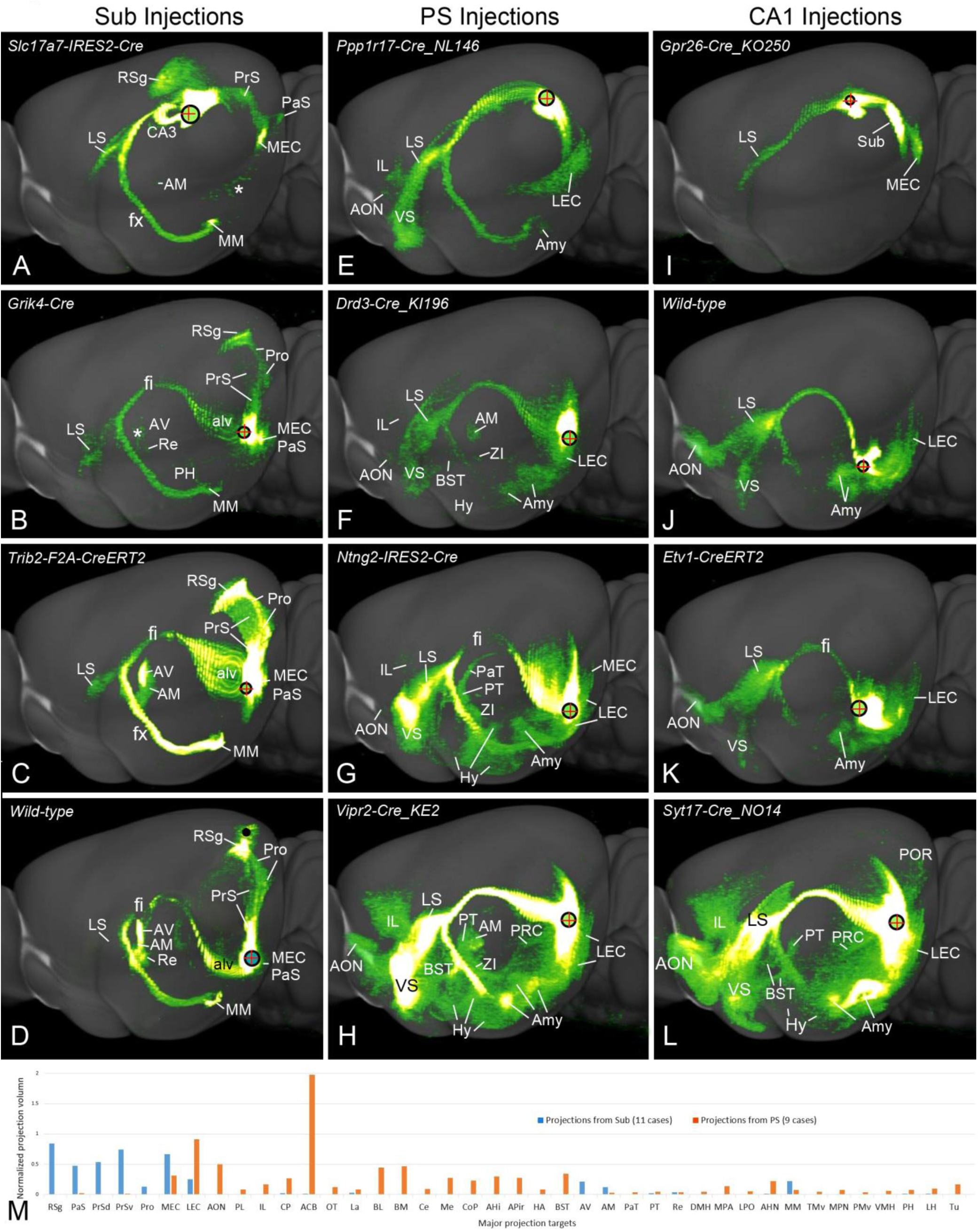
Comparison of overall projection patterns of Sub, PS and CA1. Detected fluorescent signals in each case were projected onto the lateral view of the Allen Mouse CCF template (Atlas.brain-map.org). The black circle with a red cross in each case indicates the injection site. (**A-D)** Projection patterns of the dorsal (A) and ventral (B-D) Sub in four representative cases. Axon terminal projections to RSg, PrS, PaS, MEC, AV and MM are observed in all Sub cases (see Figure 5 for detailed images of the *Trib2*-F2A-CreERT2 mouse in C). Note the dorsal injection (in A) also includes part of the underlying DG which results in terminal labeling in CA3. Note that some retrogradely labeled neurons were present occasionally in LEC and AV (white asterisks in A and B). The black dot in D indicates some injection contamination in V1. (**E-H)** Projection patterns of the dorsal (E) and ventral (F-H) PS in four representative cases. Axon terminal projections to LEC, IL, VS, Amy and AON are seen in all PS cases while those to BST and hypothalamus (Hy) are found in all PSv cases (see Figure 6 for detailed images of a *Syt17*-Cre_NO14 mouse). (**I-L)** Projection patterns of the dorsal (I) and ventral (J-L) CA1 in four representative cases. In general, CA1 shows somewhat similar projection patterns to PS but not to Sub. (**M)** Quantitative comparison of Sub (blue bars) and PS (orange bars) projections. Note the obvious differential projection targets of Sub vs. PS. For abbreviations see Table S3.

**Figure 5:**
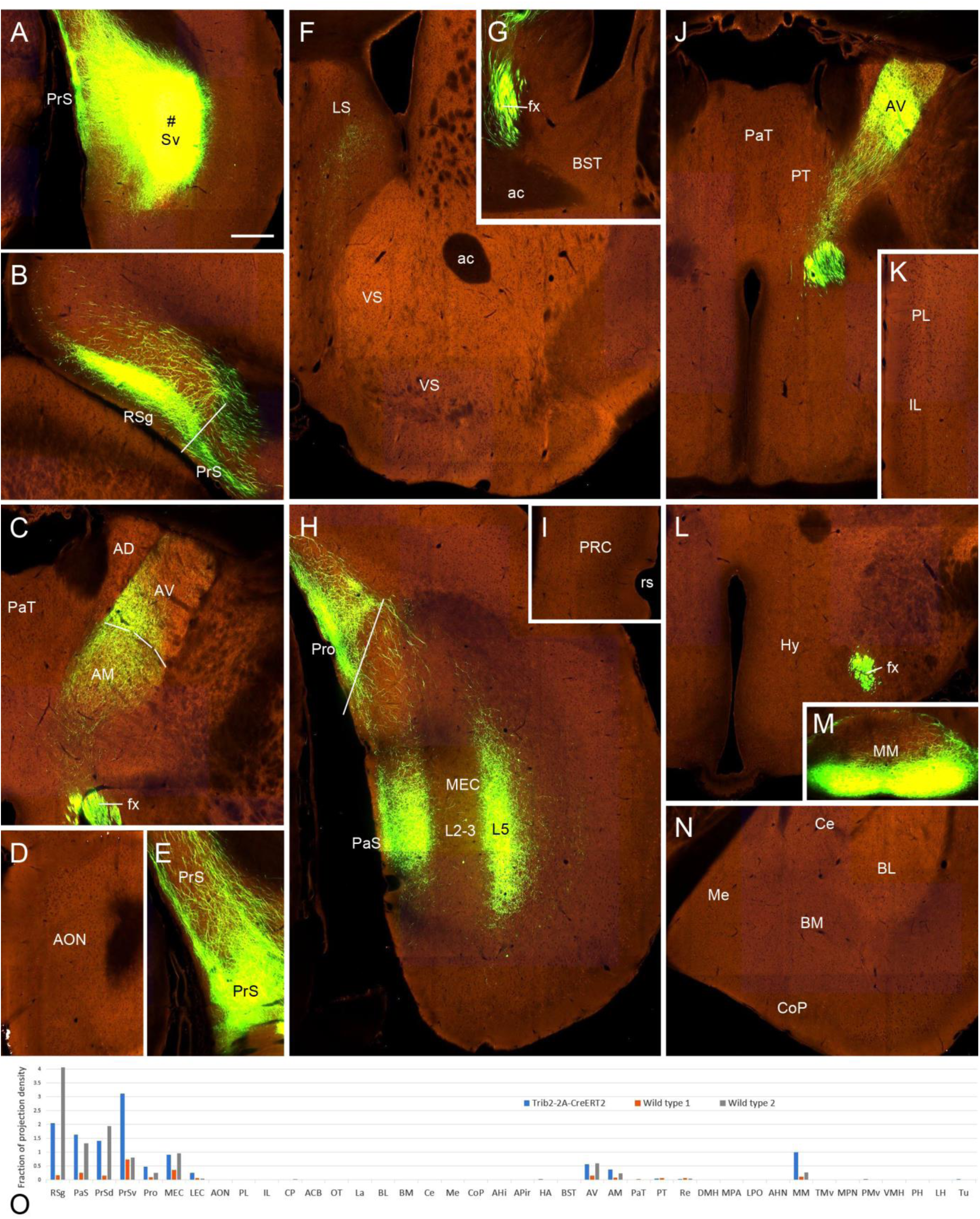
Limited efferent projections from ventral Sub (Sv). The overall projection pattern of this case was shown in Figure 4C. (**A)** A rAAV injection site in Sv (# marks the center of the injection) in a *Trib2*-F2A-CreERT2 mouse. This injection lies in the ventral Sub marked by Sv in Figure 3L, Q. (**B)** Terminal labeling in layers 2-3 of RSg. (**C)** Terminal labeling in middle AV and AM but not in AD. (**D)** Absence of terminal labeling in AON. (**E)** Fiber labeling in the dorsal PrS (The fibers pass through PrS to RSg) and strong terminal labeling in ventral PrS. (**F)** Terminal labeling in LS with no labeling in VS. (**G)** Absence of labeling in BST; (**H)** Terminal labeling in area prostriata (Pro), ventral PaS, and ventromedial MEC (mostly in layer 5); (**I)** Absence of labeling in PRC. (**J)** Terminal labeling in the anterior AV with no labeling in PaT, PT and supraoptic hypothalamic regions. (**K)** Absence of labeling in PL and IL. (**L)** Absence of labeling in the tuberal region (DMH and VMH) of the hypothalamus. (**M)** Terminal labeling in the ventral portion of MM. (**N)** Absence of labeling in the amygdala. (**O)** Quantitative comparison of Sub projections in this case (*Trib2*-F2A-CreERT2, blue bars) and in two wild-type cases (Wild types 1 and 2, orange and grey bars). Note the similar projection patterns of these cases and limited number of target regions from Sub. Bar: 280µm in A (for A-N). For abbreviations see Table S3.

A total of 12 PS-injected cases were selected and analyzed in this study. The injection sites in these cases were localized in PSd and PSv (but not in Sub) with or without involvement in other adjoining regions (Table S2). In 3 PSd cases, labeled axon terminals was clearly seen in IL, LEC, VS, LS, AON, and amygdala (Figure 4E) with few in PRC, Re and hypothalamus. In 9 PSv cases, strongly labeled axon terminals were observed in IL-PL, LEC, VS, LS, AON, amygdala, PRC, BST, PaT, PT, Re and hypothalamic nuclei (Table S2; Figures 4F-H, L, M; 6) with much less labeling in MEC and MM. In Re, the labeled terminals were concentrated in ventromedial portion (Figure 6K). In *Drd3*-Cre_KI196, *Vipr2*-Cre_KE2 and wild-type cases, labeled axon terminals were also observed in AM but not in AV (Figure 4F, H). In one *Syt17*-Cre_NO14 case, strong terminal labeling was seen in AOB (Figure 6F) although in other cases the labeling in AOB was very weak and sparse. In all 12 PS cases, no or few terminal labeling was detected in the main target regions of Sub (i.e. RSg, PrS, Pro and AV). Overall, the main targets of PS include IL, LEC, VS, LS, AON, PRC, BST, PaT, PT, Re, amygdala, and hypothalamic nuclei (target set B; Tables S2 and S4).

**Figure 6.**
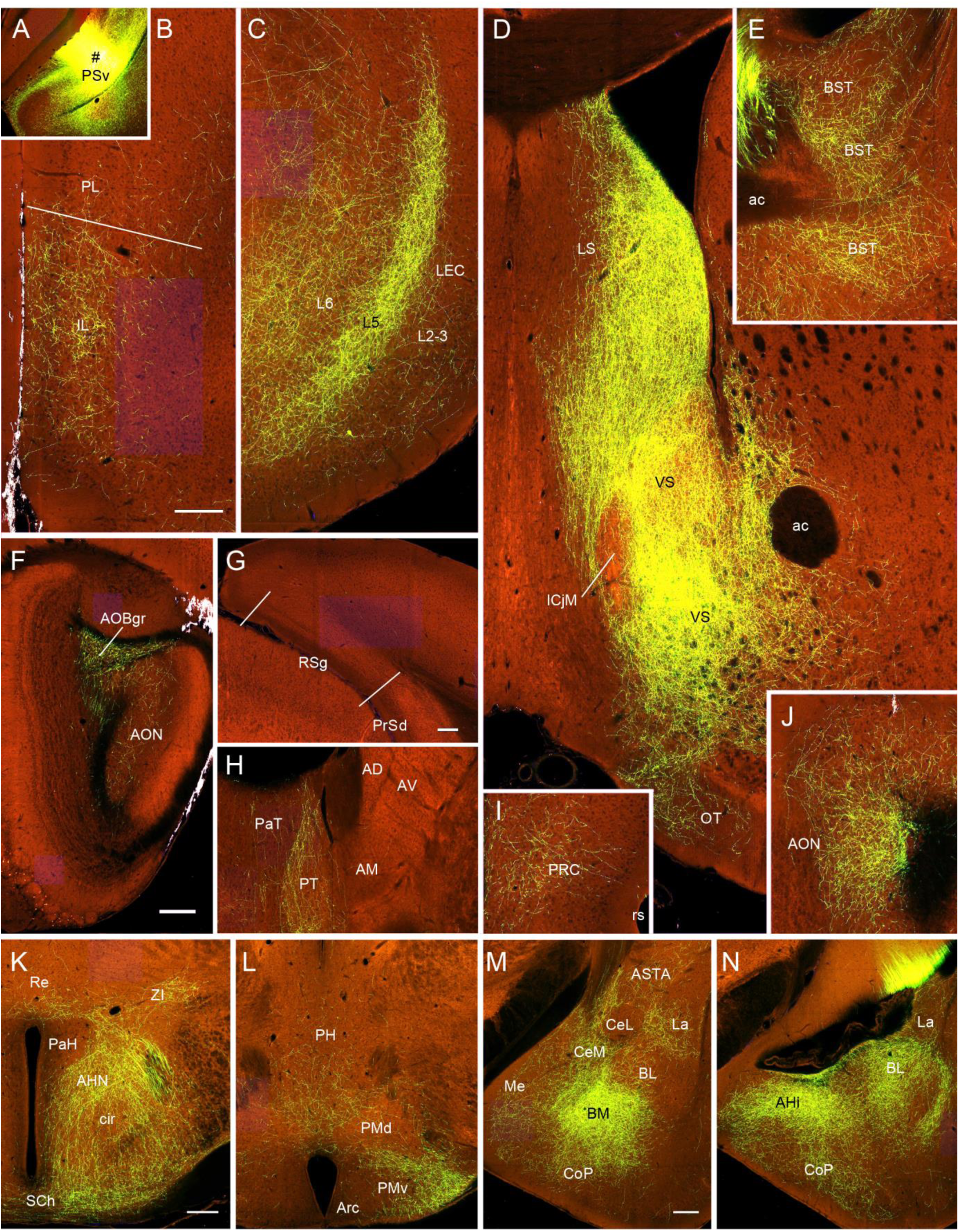
Widespread efferent projections from ventral PS (PSv). (**A**) A rAAV injection site in the ventral PS (# marks the center of the injection) of a *Syt17*-Cre_NO14 mouse. This injection lies in the most ventral PS marked by ** in Figure 3I, N. (**B-N**) Axon terminal labeling in PL, IL (B), LEC (C, mainly in layers 2-3, 5 and 6), LS, VS (D), BST (E), AOB (F, mainly in its granular layer, AOBgr), RSg, PrSd (G, no labeling), PT (H), PRC (I), AON (J), Re, ZI, AHN, SCh (K), ventral PH, lateral PMv (L), La, anterior BL, BM, Me, CeM, ASTA (M), AHi, posterior BL and CoP (N). Note much fewer labeling in major island of Island of Calleja (ICjM, D), PaT (H), PaH (K), PMd (L) and CeL (M). Note also the absence of labeling in AM (H). Overall distribution pattern of the labeled terminals in this case is similar to that in *Vipr2*-Cre_KE2 case shown in Figure 4H except strong AM labeling in the latter. Bars: 200µm in B (for B-E, I, J); 280 µm in F (for F and H); 140µm in G; 200µm in K (for K and L); 200µm in M (for M and N). For abbreviations see Table S3.

### Cell type-specific projections of Sub and PS

As demonstrated above, both Sub and PS have at least two major excitatory neuronal subclasses located in pyramidal and polymorphic cell layers, respectively. Here we explore if these two cell subclasses have different projection patterns. We first examined the Cre-mice with Sub injections and divided them into two groups: one with effective injections in both Spy and Spo and another only in Spy (Table S2). In the former group (e.g. *Trib2*-2A-CreERT2, *Grm2*-Cre_MR90 and *Slc17a6*-IRES-Cre lines; Figures 4C; S6T-W), labeled axon terminals were seen in RSg, PrS, PaS, MEC, MM and Pro (target set A1) as well as in AV, AM and Re (target set A2). In the latter group, however, the labeled axon terminals were only seen in target set A1 (e.g. *Grik4*-Cre line, Figure 4B), indicating that target set A2 is mainly innervated by Spo rather than Spy. Consistently, in cases with injections mostly restricted in Spo, labeled axon terminals were mostly observed in target set A2 (e.g. *Plxnd1*-Cre_OG1 and *Drd1a*-Cre_EY262 lines; Figure S6R, S; Table S2). Interestingly, the most distal portion of Spo (close to PrS) appears to project to AV but not AM, indicating that AM is mainly innervated by more proximal part of Spo. In fact, when an injection was restricted in the most distal portion of Spo in Sd of a *Slc17a7*-IRES-Cre mouse, labeled axon terminals were only seen in AV and not in AM (Figure S6Y). These are coincidental with our transcriptomic finding that Spo contains two clusters (RC25-26), which might innervate AV and AM, respectively. By comparing terminal labeling in Re of *Trib2*-2A-CreERT2 and *Slc17a7*-IRES-Cre mice, it is obvious to find that Re receives inputs from *Slc17a7*-Cre but not *Trib2*-Cre neurons. The Re inputs appear to derive mainly from Spo with less from Spy since strong terminal labeling was observed in Re of *Grm2*-Cre_MR90 mouse, in which *Grm2* is predominantly expressed in Spo (Figure S6E, T). Consistently, much weaker terminal labeling was observed in Re of *Grik4*-Cre and *Scnn1a*-Tg3-Cre mice, in which *Scnn1a* is predominantly expressed in Spy with less in Spo (e.g. Figure S6F, U).

We also found that PSpy and PSpo have different projections patterns. Only injections contained in PSpo resulted in terminal labeling in the thalamic nuclei AM and Re although PSpy projects to a wide range of brain regions (see above section). For example, no labeled axon terminals were found in AM and Re in *Syt17*-Cre_NO14, *Ntng2*-IRES2-Cre, *Calb1*-T2A-dgCre and *Ppp1r17*-Cre_NL146 mice (e.g. Figure 4E, G, L), in which the gene driving Cre is not expressed in PSpo (Figures 1D; 2S). On the other hand, clear projections to AM and Re were observed in *Drd3*-Cre_KI186 and *Vipr2*-Cre_KE2 mice (Figure 4F, H), in which the gene driving Cre is expressed in PSpo (Figure S6G, H). Another interesting finding was that PS (both PSpy and PSpo) did not project to AV, which instead received strong inputs from Spo. It is also worth mentioning that axon projections to PaT-PT were detected in *Syt17*-Cre_NO14, *Ntng2*-IRES2-Cre, *Calb1*-T2A-dgCre (PSpo negative; Figure 4G, L) but not in *Slc17a6*-IRES-Cre (PSpo positive; Figure S6X) mice, indicating that these projections likely originate from PSpy with no or few projections from PSpo.

We used *Slc17a6*-IRES-Cre (expression in PSpy-de), *Calb1*-T2A-dgCre (expression mainly in PSpy-su) and *Ntng2-IRES2-Cre* lines to compare the projection patterns of these cell types in PSpy. In *Slc17a6*-IRES-Cre mice with injections involving both Sub and PS, heavy projections to LS, VS and anterior hypothalamic region were detected (Figure S6X), in addition to the projections to major Sub targets (i.e. target set A: RSg, PrS, PaS, Pro, MEC, AV, AM, Re and MM). These results indicate that PSpy-de gives rise to strong projections to LS, VS and anterior hypothalamus since injections restricted in Spy of *Slc17a6*-IRES-Cre mice resulted in no or few terminal labeling in those three targets (Figure S6V, W). Interestingly, in all *Slc17a6*-IRES-Cre mice with injections contained in PSpy-de, no or few terminal labeling was observed in the amygdala (Fig. S7A-C). This indicates that PSpy-de neurons expressing Slc17a6 do not project to amygdala. In contrast, in *Calb1*-T2A-dgCre mice, heavy projections to LS, VS and amygdaloid nuclei (mainly BL) were observed with no or few labeling in hypothalamus (Fig. S7D-F; Table S2), suggesting some PSpy projections to amygdala originate from *Calb1* expressing neurons in PSpy-su. In *Ntng2*-IRES2-Cre mice, where the gene driving Cre is expressed in both PSpy-su and PSpy-de, dense axon terminals were found in all PS target regions (i.e., target set B) including BL, BM and many other amygdaloid nuclei (Fig. S7G-I; Table S2). This suggests that Ntng2 expressing neurons in PSpy-de project to BM while those in PSpy-su project to BL.

### Topographic rather than differential projections of the Sub along DV axis

With reliably defined Sub boundaries we next aim to examine and compare the main targets of Sub to clarify if topographic and/or differential projections of Sub exist along DV axis.

rAAV injections restricted in either Sd or Sv resulted in axon terminal labeling in essentially the same set of target regions (i.e., target set A: MEC, RSg, PrS, PaS, LS, AV-AM, and MM). However, the terminals were differentially distributed in their target regions. For example, labeled axon terminals from Sd were detected in the most rostrodorsal part of RSg, the most dorsal part of LS, PrS and PaS, the most dorsorostral part of MM and the most dorsolateral part of MEC (Figure S5A1-I1) as well as in the most caudolateral AM-AV (Figure S8C0-C4). Labeled terminals from Sv were observed in the most caudoventral part of Rsg, the most ventral part of LS, PrS and PaS, the most ventrocaudal part of MM and the most ventromedial part of MEC (Figure S5A2-I2) as well as in the most rostromedial AM-AV (Figure S8E0-E4). When the injections were placed in intermediate portion of Sub, labeled axon terminals were distributed in the regions between those derived from the most dorsal and most ventral parts of Sub (Figure S8D0-D4). In brief, Sub displays topographic rather than differential projections to their major targets along DV axis (i.e. target set A; Table S4), with the exception of Re where the axon terminals from both Sd and Sv appear to converge (Figure S5J1, J2). In Bienkowski et al. (2018), dorsal and ventral Sub were reported to project to target sets A and B, respectively.

### Differential and topographic projections of PS along DV axis

Since differential projections of Sub along the DV axis were not observed, we hypothesize that the DV difference of the projections from loosely defined “Subiculum” reported in literature probably originated from PS. Thus, we compared terminal distribution in the main target regions of PSd and PSv (Figures 4 and S8). PSv heavily projects to LEC, LS, VS, the amygdala and many hypothalamic nuclei; moderately to PRC, IL, AM, PT, PaT, AON, MM, and BST. In contrast, terminal labeling originated from PSd was much less dense in above-mentioned PS target regions such as LS, VS, LEC, and amygdala, and almost absent in other target regions such as IL, BST, AON, MM and hypothalamus (Table S2).

In addition to differential DV projections, topographic projections were also observed from PS to some target regions such as AM, IL, LS, VS, and BL of the amygdala. Specifically, PSd projects to caudolateral AM (Figure S8A0-A4), ventral IL, dorsomedial LS, lateroventral VS (Figure S9A-D), and lateral BL (Figure S9I-K). In contrast, PSv projects to rostromedial AM (Figure S8B0-B4), dorsal IL, ventrolateral LS, mediodorsal VS (Figure S9E-H), and medial BL (Figure S9L-N).

### Efferent projections of adjoining CA1 and MEC

Given that PS is next to CA1, we next examined the efferent projections of CA1 compared to those of PS and Sub. CA1d projects strongly to Sd, MEC and dorsal LS, lightly to PSd but not to LEC, VS and the amygdala (Figure 4I). However, CA1v projects strongly to Sv, PSv, MEC and ventral LS with weak projections to IL, PRC and caudal hypothalamus as well as to LEC, IL, AON, VS and the amygdala (Figure 4J, K; all are also the targets of PSv). In cases with injections also including adjoining PSv (e.g. Figure 4L), heavy terminal labeling was observed in IL, AON, VS, LS, LEC, and the amygdala with moderate labeling in BST, PRC, PT and caudal hypothalamus. In all cases, no labeling was found in target regions of Sub such as RSg, Pro, PrS, PaS, MM, and AV. Therefore, CA1v displays a similar projection pattern but with much less density and intensity when compared to PSv (Figure 4I-L; Table S2). In addition, CA1v projections to the amygdala mainly target BL with few to other amygdaloid nuclei.

As shown in Figure S4, Sv adjoins MEC caudally via a thin layer of white matter. To determine if MEC projects to some Sub or PS target regions we examined 6 cases with injections restricted in MEC. In two *Cux2*-IRES-Cre and two *Slc17a6*-IRES-Cre cases in which the gene driving Cre expression was restricted to layers 2-3 of MEC, the injections contained only MEC (but not Sv) and resulted in heavily labeled terminals in the molecular layer of LEC, DG, CA1, Sv and PaS with light labeling in AON, IL and OT. In 3 wild-type cases (Table S2), however, the injections also contained layers 5-6 of MEC and resulted in heavy terminal labeling in postrhinal cortex, caudate putamen, amygdala (mainly BL and Ahi), amygdalo-striatal transition area, AV, LD, NLOT, VS, and claustrum with lighter labeling in the hypothalamus. Therefore, previously reported Sv projections to the amygdala, LD, VS and hypothalamus may instead be derived from MEC that may be contained in Sub injections.

### Brain-wide differential afferent projections to Sub and PS

Since Sub and PS have differential efferent projection targets, we next examined whether Sub and PS receive differential afferent inputs. Amygdala projects to PSv with no labeling in Sv (Figure S10A-D). Dorsal and ventral CA3 project to PSd and PSv, respectively, with no projections to Sub (Figures S7J-M; S10I, J). LEC projections mainly target PS rather than Sub [in the Sub, mainly fibers rather than axon terminals were observed (Figure S10E-H)]. Injections in Re result in strong terminal labeling in PSv with much fewer labeling in Sub (Figure S10K, L). In contrast to LEC injections (Figure S10G, H), ventral MEC injections resulted in heavy terminal labeling in Sv with much fewer labeling in PSv (Figure S10M, N); in PSv, mostly fibers rather than terminals were observed. Interestingly, dorsal MEC tends to project to Sd with no labeling in Sv (Figure S10P). Finally, AV of the thalamus projects to Sub rather than PS and the labeled terminals are distributed in both dorsal Spo (Figure S10O) and ventral Spo (not shown). Quantitative analysis reveals that Sub receives its inputs mainly from MEC, PrS and AV, in addition to heavy CA1 inputs, while PS mainly receives its inputs from amygdala, LEC, CA3, Pir and Re, in addition to heavy CA1 inputs (Figure S10V).

## Discussion

The present study, for the first time, has identified 27 transcriptomic clusters/cell types in the “Sub” regions which include Sub, PS and HA. Using gene markers for specific cell types in these three regions we have accurately delineated the borders between them along full DV axis and found for the first time that the dorsal and ventral “Sub” regions are occupied mainly by Sub and PS, respectively (see Figures 7A-C and S11A-D for summary). These findings are critical to accurate and consistent localization of neurons and injections as well as data interpretation. With these findings we have demonstrated that both dorsal and ventral Sub project to the same target regions with topographic organization. This challenges traditional concept that dorsal and ventral Sub have differential projections. Furthermore, we have found the most ventral “Sub” region actually belongs to ventral PS and displays strong connections with structures critical to motivation, emotion, reward, stress, anxiety and fear. The distinction and DV difference in sizes of Sub and PS along DV axis also enable consistent interpretation of anatomic, molecular, functional and behavioral results in the literature and in different species.

**Figure 7.**
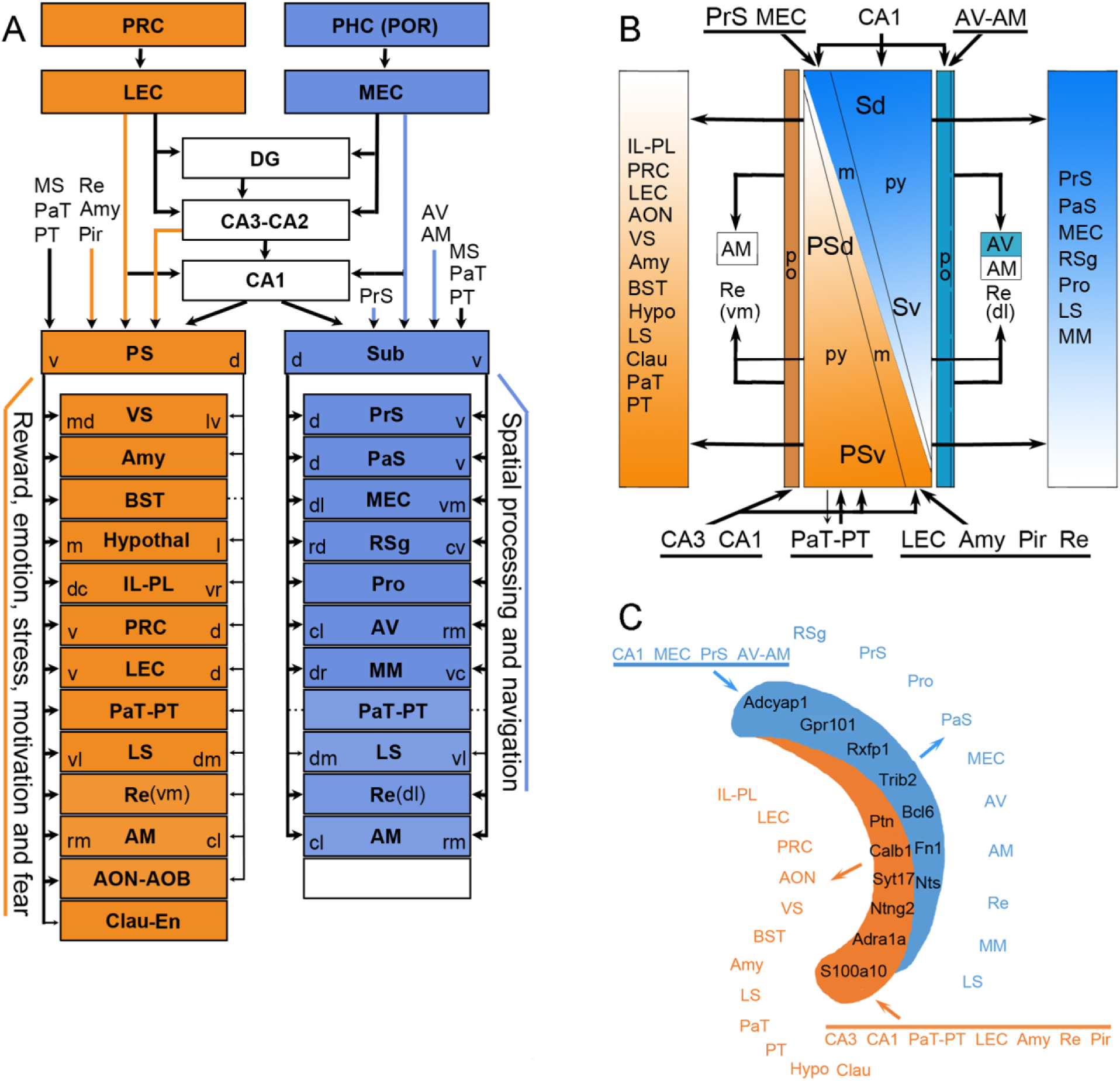
Summary of distinct circuits and transcriptomic cell types of Sub and PS. (**A**) Differential inputs and outputs of Sub and PS in the context of the entire hippocampal circuits. Sub mostly projects to regions important for spatial processing and navigation while PS mostly to widespread brain regions critical for reward, emotion, stress, motivation, anxiety and fear. Note that the PS but not Sub projections show obvious dorsoventral (DV) difference and that both PS and Sub display topographic projections to most of their targets. Potential convergence of Sub and PS projections may exist in AM, RE, MM and LS. The Hypothal indicates most of the hypothalamic regions including MM. Clau-En, claustrum and endopiriform nucleus. (**B**) Laminar organization of the inputs and outputs of Sub and PS with relation to cell types. PrS and MEC project to the molecular layer (m) while AV-AM of the anterior thalamus to the polymorphic layer (po) of Sub. CA1 projects to all layers of Sub. In PS, inputs from LEC, Amy, Pir and Re mainly project to the molecular layer while those from PaT-PT of the thalamus mainly innervate the pyramidal layer (py). Inputs from hippocampal CA3 and CA1 project to all layers. As for outputs, the polymorphic layer of Sub and PS projects to differential subdomains of the thalamic nuclei (AV, AM and Re) while the pyramidal layer of each innervates distinct cortical and subcortical targets. (**C)** Simplified summary of the molecular markers and connectivity of Sub and PS along DV axis. Note the complementary gradient in size of Sub and PS along DV axis: Sub dominates the dorsal portion while PS dominates the ventral portion of the hippocampus.

In addition to these major findings, other new findings of the present study include (Figure 7A, B): 1. PS projects to claustrum/endopiriform nucleus and accessory olfactory bulb; 2. Sub but not PS projects to area prostriata (Pro; see Lu et al., 2020), which is heavily involved in spatial processing; 3. Sub projects to both AV and AM while PS only to AM; 4. Sub projects to dorsolateral Re while PS to ventromedial Re; 5. CA3 projects to PS but not to Sub; 6. Many projection patterns of different cell types of Sub and PS using many different Cre-lines.

### Transcriptomic cell types of excitatory neurons in Sub and PS

Cembrowski et al. (2018b) recently explored the transcriptomic cell types of excitatory neurons in the subiculum regions using scRNA-seq. With a relatively small sample size (n = 1150 cells), they observed 8 clusters (with an additional one in CA1: C7). In comparison, our study revealed 27 clusters with a sample size of 8648 cells from the same subicular regions. Based on our confusion matrix analysis and the marker genes, the 9 clusters in the previous study can be mapped and partially overlap with the overall 29 clusters revealed in this study (see Figure S2A and Table S1). In this study, 9, 14 and 4 clusters or cell types are found within Sub proper, PS and HA, respectively. PSpy neurons are found to be highly heterogeneous. Cells in PSpy-su1 (the most superficial portion) are closer (i.e. more similar at transcriptional level) to the neurons in HApy, and PSpy-su2 closer to PSpy-de. Both PSpy-su2 and PSpy-de (but not Spy) are closer to CA1py. Overall, at transcriptional level, PSpy is closer to CA1py rather than to Spy, supporting our conclusion that Sub and PS are two distinct entities with differential molecular architecture, neural circuits and functional correlation (see below for further discussion). In summary, our systematic and refined cell-type identification of Sub and PS could serve as the base for accurate targeting and manipulation of specific subsets of the circuits in future studies.

### Molecular dissection of Sub and PS and their borders

Although monkey PS could be clearly identified with help of AChE stain by many researchers (Barbas and Blatt, 1995; Blatt and Rosene, 1998; Ding, 2013; Fudge et al., 2012; Rosene and van Hoesen, 1987; Sounders et al, 1988b; Yukie, 2000), reliable and precise identification of rodent PS has proven to be very difficult due to the curvature of the hippocampus and the lack of reliable markers. Consequently, the PS region has been treated as part of Sub in most rodent literature. However, the existence of distinct rodent PS has been re-emphasized recently based on a combined analysis of comparative, connectional, neurochemical and molecular data (Ding, 2013). In the present study, we have registered unbiased transcriptional cell type classification to anatomical regions and found that the general “subiculum” contains at least three distinct regions: PS, Sub proper and HA. Furthermore, the borders, extent and topography of Sub, PS and HA along DV axis are consistently and reliably defined based on distinct molecular markers revealed from the transcriptome. We have further uncovered that the sizes (widths) of Sub and PS decrease and increase respectively along DV levels (see Figures 3, S4 and 7C) and that Sub and PS display generally distinct afferent and efferent projections (Figure 7A, B).

Based on these findings, many confusing and conflicting results in previous rodent studies can be explained with the introduction of the PS concept and the precise and reliable demarcation of its borders. For example, we found that the region previously labeled as ventral Sub in most rodent literature in fact is the ventral PS because it expresses transcriptionally identified marker genes for PS rather than those for Sub (see Figures 3 and S4; Table S1). Consistent with this conclusion, this region, although called “ventral Sub”, was found to project to IL, VS, AON, BST, PRC and many amygdaloid and hypothalamic nuclei (Bienkowski et al., 2018; Cullinan et al., 1993; Kishi et al., 2006; McDonald, 1998; Swanson and Cowan, 1977), all of which are typical target regions of PS revealed in the present study. Thus, previously reported projections from “ventral Sub” to above target regions mostly originate from ventral PS. Since these target regions are strongly associated with functions such as motivation, emotion, reward, stress, anxiety and fear (Aggleton 2012; Andrzejewski et al, 2006; Herman and Mueller, 2006; O’Mara et al., 2009; Potvin et al., 2006; Subhadeep et al., 2017), We conclude here that it is PS rather than Sub predominantly projecting to the structures critical for motivation, emotion, reward, stress, anxiety and fear.

Many previous studies in rodent divided the “Sub” (mainly the dorsal “Sub”) into distal and proximal “Sub” instead of Sub and PS (see Figure S11 for summary, and Aggleton, 2012; Aggleton and Christiansen, 2015; Witter, 2006 for reviews). The distal and proximal “Sub” were originally used to describe the locations of injection sites and labeled neurons in the dorsal “Sub” (e.g. Naber and Witter, 1998; Witter, 2006; Witter et al., 1990). Thus, the “Sub” region closer to CA1 was named proximal “Sub” while that away from CA1 named distal “Sub” with no specific markers used to demarcate the borders. Recently selected gene markers were introduced to mark the distal and proximal “Sub” (S-dis and S-pro, respectively) (Cembrowski et al., 2018a, 2018b) or Sub and PS (Bienkowski et al., 2018; Ding, 2013), mostly for dorsal “Sub”. Connectivity data appear to support this subdivision of the dorsal “Sub” (e.g. Bienkowski et al., 2018; Cembrowski et al., 2018a; Witter, 2006). However, in the ventral part of the “Sub” region, distal and proximal “Sub” do not appear to be dividable (e.g. Cembrowski et al., 2018b) or could not be divided into S-dis and S-pro (Bienkowski et al., 2018; see Figure S11). In fact, as shown in Figure S11, the distal and proximal “Sub” express Sub and PS genes such as *Nts* and *Ntng2*, respectively, in the dorsal part. In the ventral part, however, both distal (away from CA1) and proximal (close to CA1) regions express PS genes (e.g. *Ntng2* and *S100a10*). At more caudal levels, it is even harder to demarcate distal and proximal “Sub” (see Figure S11). Therefore, the concept of distal and proximal “Sub” cannot be clearly applied to the ventral subicular regions. Here, using distinct markers consistently defined for Sub and PS cell types from our transcriptomic taxonomy, we have revealed that the ventral “Sub” region actually belongs to ventral PS (Figures 3, 7C, S4, S11) rather than ventral Sub.

### Untangling of the wiring circuits of Sub and PS

Guided with reliable boundaries of Sub and PS along the DV axis (Figures 3, S4), we are able to untangle the wiring circuits of Sub and PS. When the anterograde tracers were restricted in Sub, labeled axon terminals were mostly observed in RSg, PrS, PaS, MEC, Pro, LS, Re (dl), AV, AM, and MM (i.e. target set A). In contrast, when the tracers were strictly placed in PS, the terminal labeling was found in IL, LEC, VS, LS, Re (vm), AON, BST, PRC, PaT-PT, claustrum, amygdala, and almost all hypothalamic regions (i.e. target set B; see summary in Figure 7A, B). Thus, Sub and PS basically project to distinct sets of brain regions with only a few exceptions. Three possible exceptions are in LS, AM and MM where both Sub and PS projections may converge. This is in contrast to previous studies showing mixed (not distinct) projection patterns of the “Sub”. The reasons for the previous mixed projection patterns may be four fold. The first is due to the mixing of two different entities (PS and Sub) into one “Sub”. The second is due to the oblique border between the two entities and the small size of each. It is thus very difficult to restrict tracer injections into one entity without leaking into the other in conventional tracing experiments. Third, anterograde tracer injections targeting the most caudal Sub could leak into adjoining MEC. As reported in this study, MEC also has strong projections to amygdala, and thus could be interpreted as originating from Sub. Fourth, Sub injections in wild-type mice could reach underlying white matter where axon fibers of passage from PS could take up the tracers and result in some labeling in PS target regions. Cre-dependent viral tracing used in this study could increase the accuracy of tracer injections (e.g. Figure S6).

The finding of distinct projection patterns of Sub and PS is consistent with previous retrograde tracing results from the dorsal distal and proximal ‘Sub”. For example, when retrograde tracers were injected in IL, VS, BST, amygdala and PRC, labeled neurons were found only in the proximal “Sub” region corresponding to PS in this study (Christie et al, 1987; Ishizuka, 2001; Kishi et al., 2006; Ottersen, 1982; Phillipson and Griffiths, 1985; Shi and Cassell, 1999; Veening, 1978; Weller and Smith, 1982; Witter, 2006; Witter et al., 1990). When the retrograde tracers were injected into RSg and AV, labeled neurons were only observed in the distal “Sub” corresponding to the Sub proper revealed in the present study (Christiansen et al., 2016; Meiback and Siegel, 1977b; Wyss, and Van Groen, 1992). However, the ventral ‘Sub” region could not be subdivided into distal (away from CA1) and proximal (close to CA1) parts and it actually belongs to ventral PS, as demonstrated in the present study. Thus, distinguishing Sub from PS consistently along DV axis is very helpful in interpreting inconsistent and confusion results from previous studies. Consistently, afferent projections to Sub and PS were found to originate generally from differential brain regions. As summarized in Figure 7A-B, PS receives major inputs from CA3, CA1, LEC, piriform cortex, midline thalamic nuclei, medial septal nucleus and amygdala while Sub receives its inputs mainly from CA1, MEC, PrS, and AV-AM (also see Agster and Burwell, 2013; Amaral et al., 1991; Ding, 2013; Roy et al., 2017; Tamamaki et al., 1987).

Taken together, the Sub and PS defined in this study apply well to both dorsal and ventral “Sub” regions and display distinct wiring circuits with PS having more widespread inputs and outputs than Sub. The Sub connects with regions heavily involved in the procession of spatial information and navigation. In contrast, PS connects with limited cortical regions but many subcortical regions that have strong association with motivation, reward, emotion, fear and stress.

### Topographic and differential projections of Sub and PS along DV axis

With the exception of Pro and Re, all other main target regions of Sub receive clear topographic projections from Sub. These findings are consistent with previous retrograde tracing results in rat (Allen and Hopkins, 1989; Honda and Ishizuka, 2015; Meibach and Siegel, 1977a; Wyss and Van Groen, 1992) and mouse (Cembrowski et al., 2018a) because retrogradely labeled neurons in these studies are actually restricted in the region corresponding to the Sub proper of the present study. Moreover, the dorsal and ventral limits of Sub are highly consistent with the dorsal and ventral borders revealed with the molecular markers in the present study. Thus, the topographic projections from Sub to its main target regions such as RSg, MEC, and PrS could also be used to confirm the most dorsal and the most ventral limits of Sub, as shown in Figure S4. Another important finding is the absence of DV difference of Sub projections because Sub at all DV levels have dense projections to RSg, MM, MEC, PrS, PaS and AV-AM.

In previous studies, however, connectional difference of the “Sub” region and CA1 along DV axis was frequently reported (see Strange et al., 2014). In this study, we found this DV difference exists for PS and CA1 projections but not for Sub projections. The main reason for this discrepancy is that many previous studies did not recognize PS but treated it as part of the “Sub” (Canteras and Swanson, 1992; Cullinan et al, 1993; Swanson and Cowan, 1977) and/or did not use reliable markers to distinguish Sub from PS. Using reliable molecular markers in this study we found that PS and adjoining CA1 have obvious differential DV projections. This difference includes (1) ventral PS has dense while dorsal PS has no or few projections to AON, BST and hypothalamus; (2) ventral PS has much stronger projections to VS, LS, and amygdala than dorsal PS does. Interestingly, PS projections to VS, IL, LS, and BL of the amygdala show rough topographic organization.

Another important finding of the present study is that Sub and PS dominates the dorsal and ventral “subicular” region, respectively, along DV axis (see Figures 3, S4 and 7C). In fact, we found the most dorsal region lacks PS (Figures 3A, S4A, B) while the most ventral region lacks Sub (Figures 3H-J, N, O; S4F, G, N, O), and the size and extent of Sub and PS display opposite DV gradients (Figure 7C). This finding indicates that dorsal “subicular” lesion mainly damages Sub while ventral “subicular” lesion mainly damages PS. This result explains well the previous lesion and behavioral findings that dorsal lesion mainly displayed impaired spatial processing and navigation while ventral lesion mainly showed impaired reward, emotion, motivation, fear and stress response (O’Mara et al., 2009; Strange et al., 2014).

### Functional implications

Consistent with many previous works, the present study demonstrated that the main targets of Sub projections include RS (RSg), MEC, PrS, PaS, MM, ATN (AV and AM) and Re. Lesions in these structures and in the dorsal “Sub” heavily impair spatial memory, orientation and navigation (Aggleton and Christiansen, 2015; Cembrowski et al, 2018a; Potvin et al, 2009). Physiologically, these structures contain cells sensitive to spatial information such as grid cells, head direction cells, boundary vector cells and cells encoding animal’s current axis of travel relative to environmental boundaries (Hafting et al, 2005; Jankowski et al, 2014; Lever et al, 2009; Olson et al, 2017; Taube, 2007). The topographic organization of Sub projections to the main target structures (rat: Honda and Ishizuka, 2015; Wyss and Van Groen, 1992; mouse: this study) and gradient gene expression along DV axis (Strange et al, 2014) of the hippocampus including Sub also appear to support spatial processing and computation.

Many previous studies reported ventral “Sub” projections to IL, LEC, PRC, LS, VS, BST, AON, Pat-PT, amygdala and hypothalamus (see Bienkowski et al., 2018; Ding, 2013; Jin et al., 2015). Lesions or simulations in these structures and in the ventral “Sub” resulted in changed feeding and social behavior and stress responses (Aqrabawi et al., 2016; Cassel et al., 2013; Farrell et al., 2010; Hsu et al., 2014; Mannella et al., 2013; Parfitt et al., 2017; Riaz et al. 2017; Sweeney et al. 2015, 2016; Vranjkovic et al. 2017; Wassum and Izquierto, 2015). In this study we recognize that this ventral “Sub” actually belongs to ventral PS in terms of its gene expression and connectivity patterns. Recently, the “CA1v” region, corresponding to PSv and adjoining CA1v of the present study, has been found to contain a lot of “anxiety” cells which mainly project to hypothalamus and drive anxiety-related avoidance behavior and aversion (Jimenez et al, 2018). This same region also contains another set of cells that project to amygdala and mainly modulate contextual fear memory encoding and retrieval (Jimenez et al, 2018). One striking finding of this study is that PS contains at least 14 clusters (cell types). We hypothesize that different cell types might innervate a subset of target regions of PS and thus modulate a specific subset of neurons responsible for different function in fear, anxiety, reward, motivation, stress and addiction. For example, *Slc17a6* expressing neurons in PSpy-de do not appear to innervate amygdala while *Calb1* expressing neurons in PSpy-su do (Fig. S7). Therefore, *Calb1*-Cre mice could be used to specify the function of the PS projections to amygdala in future studies.

In summary, as demonstrated in this study, Sub and PS are two distinct regions with differential transcriptome, molecular signature, connectivity and functional correlation along the entire DV axis. The benefits of introducing/rescuing the term PS for rodent brain include at least the following aspects: 1. Enable consistent description of Sub and PS across species; 2. Match well to distinct molecular markers and connectivity of Sub and PS along DV axis; 3. Correlate well with differential functions of Sub and PS; 4. Enable accurate targeting and description for future lesion, injection, stimulation and recording studies; 5. Facilitate accurate evaluation and interpretation of the results from animal models of related diseases.

## STAR*METHODS

All experimental procedures were approved by the Allen Institute Animal Care and Use Committee and conform to NIH guidelines.

### Single-cell isolation for SMART-Seq

We adapted a previously described protocol to isolate neurons from the mouse brain (Tasic et al., 2018). Briefly, adult male and female mice (P56 ± 3; n=4) from the pan-glutamatergic mouse line *Slc17a7-IRES2-Cre;Ai14* were anesthetized with isoflurane and perfused with cold carbogen-bubbled artificial cerebrospinal fluid (ACSF). The brain was dissected, submerged in ACSF, embedded in 2% agarose, and sliced into 250-μm coronal sections on a compresstome (Precisionary). Under microscope, and with reference to Allen Mouse Brain CCF (v3), full DV extent of the regions PS-Sub-HA and PrS-PoS-PaS were microdissected from the 250-μm thick slices with a knife and dissociated into single cells with 1 mg/ml pronase (Sigma, Cat#P6911-1G). Single cells were isolated by FACS into individual wells of 8-well PCR strips containing lysis buffer from the SMART-Seq v4 kit with RNase inhibitor (0.17 U μl−1), immediately frozen on dry ice, and stored at −80 °C.

### Single-cell isolation for 10X Chromium v2

Adult male and female mice (P56 ± 3; n=2) from the pan-neuronal mouse line *Snap25-IRES2-Cre;Ai14* were anesthetized, brains were dissected, and 250-μm thick coronal slices were prepared as described for SMART-Seq processing. The entire region containing full DV extent of PS-Sub-HA-PrS-PoS-PaS were micro-dissected out with a knife and a microscope, under which the anatomic borders can be identified, and digested with 30 U/ml papain (Worthington #PAP2) in ACSF for 30 mins at 35 °C in a dry oven, with a targeted solution temperature of 30 °C. Enzyme digestion was quenched by exchanging the papain solution three times with quenching buffer (ACSF with 1% FBS and 0.2% BSA). Samples were incubated on ice for 5 minutes before trituration. In 1 ml of quenching buffer, the tissue pieces were triturated through a fire-polished pipette, with 600-um diameter opening, approximately 20 times. Tissue pieces were allowed to settle, and the supernatant, which now contains suspended single cells, were transferred to a new tube. Fresh quenching buffer (1 ml) was added to the settled tissue pieces, and trituration and supernatant transfer was repeated using 300-um and 150-um fire polished pipettes. Final volume of the single cell suspension is 3 ml. It is possible for small tissue pieces to remain in the original tube after these three rounds of trituration, and these are discarded. To remove excessive debris, the single cell suspension was passed through a 70-um filter into a 15-ml conical tube with 500 µl of high BSA buffer (ACSF with 1% FBS and 1% BSA) at the bottom to help cushion the cells during centrifugation at 100xg in a swinging bucket centrifuge for 10 minutes. The supernatant was discarded, and the cell pellet was resuspended in 1 ml quenching buffer. Cells were passed through 70-um filter again and DAPI (2 ng/ml) was added to the suspension. Cells were isolated using FACS (BD Aria II) gated on DAPI and tdTomato. In order to increase yield while reducing the sheath volume in the sorted suspension, we sorted on Fine Tune mode which has Yield, Purity, and Phase Mask all set to 0. We divided up the sample so that we could sort 30,000 cells at a time, within 10 minutes, into 5-ml tube containing 500 µl of quenching buffer. Each aliquot of sorted 30,000 cells were immediately centrifuged at 230xg for 10 minutes in a swinging bucket centrifuge with 200 µl of high BSA buffer at the bottom for cushion. No pellet can be seen with this small number of cells, so we take out the supernatant and leave behind 35 µl of buffer, in which we resuspended the cells. The resuspended cells are stored at 4 °C until all samples have been collected for chip loading on the 10X Genomics controller. Our typical sort takes 30 minutes for three aliquots. We observe more cell death for longer sorts. Typically, one aliquot of 30,000 sorted cells result in a final suspension of 5,000 - 20,000 viable cells for loading onto one port of the 10X Genomics chip.

### cDNA amplification and library construction

We used the SMART-Seq v4 Ultra Low Input RNA Kit for Sequencing (Takara, 634894) to reverse transcribe poly(A) RNA and amplify full-length cDNA. We performed reverse transcription and cDNA amplification for 18 PCR cycles in 8-well strips, in sets of 12–24 strips at a time. All samples proceeded through Nextera XT DNA Library Preparation (Illumina FC-131-1096) using Nextera XT Index Kit V2 (FC-131-2001). Nextera XT DNA Library prep was performed according to manufacturer’s instructions except that the volumes of all reagents including cDNA input were decreased to 0.4× or 0.5× by volume. Subsampling of the reads to a median of 0.5 million per cell results in similar gene detection per cell (>89% of genes detected, data not shown), showing that we detect most of the genes at 2.5 million reads per cell. Details are available in ‘Documentation’ on the Allen Institute data portal at: http://celltypes.brain-map.org. For 10X Genomics processing, we used Chromium Single Cell 3’ Reagent Kit v2 (10X Genomics #120237). We followed manufacturer’s instructions for cell capture, barcoding, reverse transcription, cDNA amplification, and library construction. Average sequencing depth was ∼59k reads per cell across 9 libraries.

### Sequencing data processing and quality control

For the SMART-Seq V4 dataset, fifty-base-pair paired-end reads were aligned to GRCm38 (mm10) using a RefSeq annotation gff file retrieved from NCBI on 18 January 2016 (https://www.ncbi.nlm.nih.gov/genome/anno-tation_euk/all/). Sequence alignment was performed using STAR v2.5.3 with twopassMode. PCR duplicates were masked and removed using STAR option ‘bamRemoveDuplicates’. Only uniquely aligned reads were used for gene quantification. Gene read counts were quantified using the summarizeOverlaps function from R GenomicAlignments package using both intronic and exonic reads, and QC was performed as described in (Tasic 2018). The 10X dataset was processed using cellranger v3.0.0 pipeline. Doublet detection was performed using scrattch.hicat doubletFinder function, adapted from the original doubletFinder package v1.0 (https://github.com/chris-mcginnis-ucsf/DoubletFinder) for better efficiency and performance. 10X doublet cells were defined as cells with doublet score greater than 0.3, and removed before clustering. We determined 10X cell class based canonical markers into neurons and non-neuronal cells. For neuronal cells, we selected cells with at last 2000 detected genes, and for non-neuronal cells, we selected cells with at least 1000 detected genes.

### Clustering

Clustering for both SMART-Seq and 10X datasets were performed using house developed R package scrattch.hicat (available via github https://github.com/AllenInstitute/scrattch.hicat). In addition to classical single-cell clustering processing steps provided by other tools such as Seurat, this package features automatically iterative clustering by making finer and finer splits while ensuring all pairs of clusters, even at the finest level, are separable by fairly stringent differential gene expression criteria. The package also performs consensus clustering by repeating iterative clustering step on 80% subsampled set of cells 100 times, and derive the final clustering result based on cell-to-cell co-clustering probability matrix. This feature enable us to both fine tune clustering boundaries and to assess clustering uncertainty. For differential gene expression criteria between clusters, q1.th = 0.4, q.diff.th=0.7, de.score.th=150, min.cells=20 is used for 10X cells, and q1.th = 0.5, q.diff.th=0.7, de.score.th=150, min.cells=4 is used for SMART-Seq cells. Clusters for each dataset were inspected manually, and based on marker genes, clusters that believed to be outside of subicular complex were eliminated from downstream analysis.

### Consensus clustering between 10X and SMART-Seq

To provide one consensus subicular cell type taxonomy based on both 10X and SMART-Seq datasets, we developed a novel integrative clustering analysis across multiple data modalities, now available via unify function of scrattch.hicat package. Unlike Seurat CCA approach (Butler et al., 2018) and scVI (Lopez et al., 2018), which aim to find aligned common reduced dimensions across multiple datasets, this method directly builds a common adjacency graph using all cells from all datasets, then applies standard Louvain community detection approach for clustering. To build the common graph, we first chose a subset of reference datasets from all available datasets, which either provides stronger gene detection and/or more comprehensive cell type coverage. The key steps of the pipeline are outlined below:

1. **Select anchor cells for each reference dataset.** For each reference dataset, we random sampled up to 5000 cells as anchors. If independent clustering results for the reference datasets are available, we sample at least100 anchor cells per cluster to achieve more uniform coverage of cell type.
2. **Select high variance genes.** High variance genes and PCA dimensions reduction were performed using scrattch.hicat package. PCA dimensions that highly correlated with technical bias such as gene detection or mitochondria gene expression were removed. For each remaining PCA dimension, Z scores were calculated for gene loadings, and top 100 genes with absolute Z score greater than 2 were selected. The high variance genes from all references datasets were pooled.
3. **Compute K nearest neighbors.** For each cell in each dataset, we computed its K nearest neighbors among anchor cells in each reference datasets based on the high variance genes selected above. Different distance metrics can be selected for computing nearest neighbors between different pairs of datasets. By default, Euclidean distance is used when query and reference dataset is the same. Between different datasets, correlation is used as similarity metrics to select K nearest neighbors.
4. **Compute the Jaccard similarity**. For every pair of cells from all datasets, we compute their Jaccard similarity, defined as the ratio of the number of shared K nearest neighbors (among all anchors cells) over the number of combined K nearest neighbors.
5. **Perform Louvain clustering based on Jaccard similarity.**
6. **Merge clusters.** To ensure that every pair of clusters are separable by conserved differentially expressed (DE) genes across all datasets, for each cluster, we first identified the top 3 nearest clusters. For each pair of such close-related clusters, we computed the differentially expressed genes in each dataset, and choose the DE genes that are significant in at least one dataset, while also having more than 2 fold change in the same direction in all datasets. We then compute the overall statistical significance based on such conserved DE genes for each dataset independently. If any of the dataset pass our DE gene criteria (Tasic et al., 2018), the pair of clusters remained separated, otherwise they were merged. DE genes were recomputed for merged clusters, and the process repeat until all clusters are separable by sufficient number of conserved DE genes. If one cluster has fewer than the minimal number of cells in a dataset, then this dataset is not used for DE gene computation for all pairs involving the given cluster. This step allows detection of unique clusters only present in a subset of clusters.
7. Repeat steps 1-6 for cells within cluster to gain finer resolution clusters until no clusters can be found.
8. Concatenate all the clusters from all the iterative clustering steps, and perform final merging as described in step **6.**

This integrative clustering pipeline allows us to resolve clusters at fine resolution, while ensuring proper alignment between datasets by requiring presence of conserved DE genes. It also allows us to leverage the strengths of different datasets. For example, between clusters that are separated by weakly expressed genes, SMART-Seq dataset provides the statistical power for separation, and the relevant genes help to separate 10X cells into clusters with consistent fold change. On the other hand, for clusters that have very few cells in SMART-Seq, 10X provides the statistical power for separation, and relevant genes are used to split SMART-Seq cells accordingly.

We applied this pipeline using both SMART-Seq and 10X datasets as reference, and the consensus clustering results were highly concordant with independent clustering results. All the major cell type markers are highly conserved at the cluster level. We calculated conserved DE genes between all pairs of clusters, and calculated the cluster means of these genes for each dataset. The concatenated cluster mean expression profiles across all datasets are used to build cell type taxonomy tree. Using the K nearest neighbors, we imputed the gene expression of SMART-seq cells based on 10X anchor genes, and imputed gene expression is used to create tSNE plot.

### Confusion matrix analysis workflow

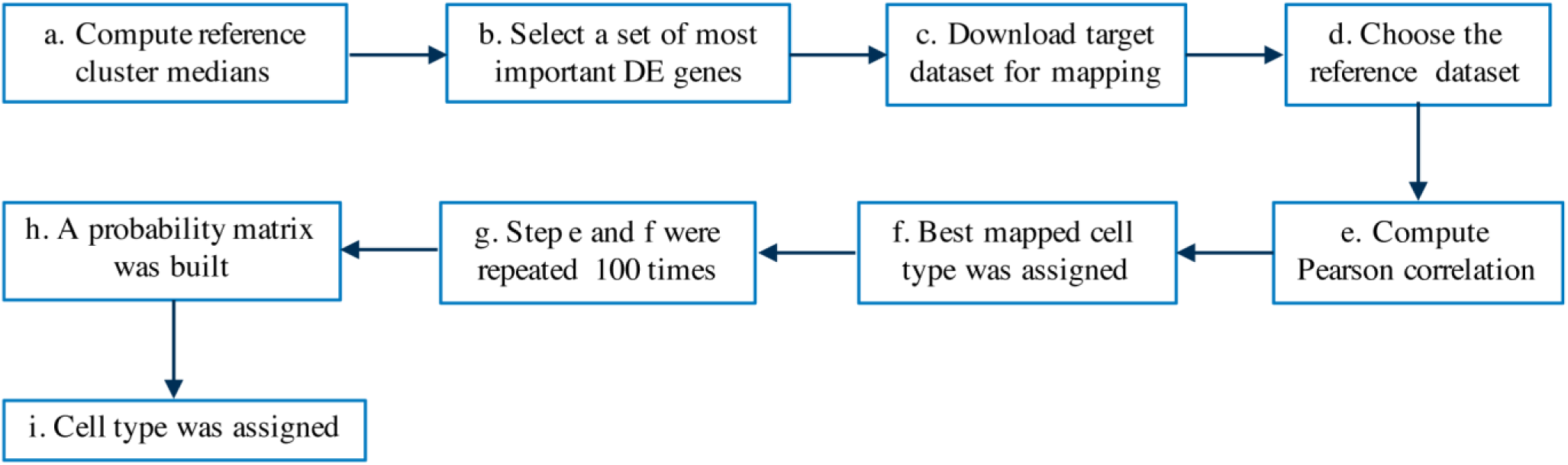
(a) Compute reference cluster medians: median gene expressions for each cell type (in this work) was computed for Smartseq cells and for 10X cells individually; (b) select a set of most important DE genes: a set of most important differentially expressed genes (∼4570) was selected (this was done during clustering); (c) download target dataset for mapping: Gene expression and the cell types (clusters) of the target dataset (e.g. dataset from Cembrowski et al., 2018b) was downloaded; (d) choose the reference dataset: depending on the target data collection method, either Smartseq or 10X reference was chosen; (e) compute Pearson correlation: Pearson correlation was computed between each target cell gene expression and the cluster median gene expression of the reference dataset. A random selection of 80% of selected genes were used; (f) best mapped cell type was assigned: reference cell type which has the highest correlation with the target cell was assigned as the best mapped cell type for that cell; (g) step e and f were repeated 100 times: each time 80% of selected genes were used at random; (h) a probability matrix was built: using the result of step g, a probability matrix was built which shows what is the probability of a target cell to be mapped to each of the reference cell types; (i) cell type was assigned: for each target cell, the cell type which has the highest mapped probability was assigned as the corresponding cell type.

### *In situ* hybridization (ISH) and anatomical mapping of the clusters

ISH data used for anatomical registration and spatial validation of the transcriptional clusters are from Allen Mouse Brain Atlas (Lein et al, 2007), which is publicly available at www.brain-map.org. Detailed description can be found at Allen Mouse Brain Atlas documentation page (http://help.brain-map.org/display/mousebrain/Documentation). Generally, 20-50 marker genes for each cluster were selected from transcriptome and their expression in Sub, PS and adjoining regions was examined and validated with Allen Mouse Brain Atlas ISH dataset. Afterwards, representative ISH images for the locations of specific clusters were downloaded and displayed as shown in Figures 2, S1 and S3 and Table S1.

### Selection of gene markers for Sub and PS

Since the “subicular” region distributes along a long DV axis, we chose the well-characterized dorsal portion for gene differentiation between Sub and PS. Based on unbiased transcriptomic clustering, we chose strongly and selectively expressed genes as marker genes for Sub and PS, respectively. For example, *Ntng2* and *Calb1* are strongly expressed in the region close to CA1 (i.e. away from PrS; see Figures 3, S4) but not in the region close to PrS (i.e. away from CA1) and these two genes could be treated as PS markers based on the concept and definition of PS (Bienkowski et al., 2018; Ding, 2013; Lorente de No, 1934; Rosene and Van Hoesen, 1987; Saunders et al., 1988a, b). Consistently, the genes expressed strongly and selectively in the region close to PrS but not in the region close to CA1 (e.g. *Nts* and *Bcl6*) were treated as Sub markers (see Figures 3, S4). These selected gene markers were then applied to the whole “subicular” region along DV axis to obtain consistent and reliable boundaries of Sub and PS, which is critical for evaluation of tracer injections.

### Mice used for tracing studies

Wild type (C57BL/6J; n=20) and Cre driver transgenic mice (n=72) at postnatal day (P) 56 ± 3 were used in tracing study. The Cre lines mainly includes Cux2-IRES-Cre (n=8), Calb1-T2A-dgCre (n=3), Dlg3-Cre_KG118 (n=1), Drd1a-Cre_EY262 (n=1), Drd3-Cre_KI196 (n=3), Etv1-CreERT2 (n=3), Grik4-Cre (n=8), Grm2-Cre_MR90 (n=1), Gpr26-Cre_KO250 (n=4), Grp-Cre-KH288 (n=3), Ntng2-IRES2-Cre (n=3), Otof-Cre (n=3), Pcdh9-Cre_NP276 (n=2), Plxnd1-Cre_OG1 (n=2), Ppp1r17-Cre_NL146 (n=5), Rorb-IRES2-Cre (n=2), Scnn1a-Tg3-Cre (n=3), Slc17a6-IRES-Cre (n=5), Slc17a7-IRES2-Cre (n=2), Syt17-Cre_NO14 (n=6), Trib2-2A-CreERT2 (n=1) and Vipr2-Cre_KE2 (n=3). These lines were generated at the Allen Institute or imported from external sources (see Harris et al, 2014) and examples of Cre expression in Sub, PS and adjoining regions from these lines were shown in Figure S6 and Table S2.

### Animal surgery and tracer injection

The methods for animal surgery and tracer injection were reported previously (Oh et al., 2014) and can be found at the Allen Mouse Brain Connectivity Atlas documentation page (http://help.brain-map.org//display/mouseconnectivity/Documentation). Briefly, both wild type and Cre mice were anesthetized with 5% isoflurane and mounted onto a stereotaxic frame (model 1900; Kopf, Tujunga, CA) prior to surgery. During surgery, anesthesia was maintained at 1.8–2% isoflurane. For subicular and prosubicular injections along the D-V axis, a glass pipette (inner tip diameter 10-20 µm) loaded with AAV was lowered to the desired depth based on the atlas of Paxinos and Franklin (2012). For wild-type mice, a pan-neuronal AAV vector expressing EGFP under the human synapsin I promoter (AAV2/1.pSynI.EGFP.WPRE.bGH) was injected in target regions, while in Cre driver mice a Cre-dependent AAV (AAV2/1.pCAG.FLEX.EGFP.WPRE.bGH) was injected. The AAV (serotype 1, produced by UPenn viral core; titer > 10^12^ GC/ml) was delivered by iontophoresis (current 3 µA and 7 seconds on/7 seconds off duty cycle) for 5 minutes. After tracer injections, the skin incision was sutured and the mice were returned to their cages for recovery.

### Brain preparation and imaging

After 21 days of survival time, mice were intracardially perfused with 10 ml 0.9% NaCl followed by 50 ml freshly prepared 4% paraformaldehyde (PFA) after anesthetization with 5% isoflurane. After extraction, brains were postfixed in 4% PFA at room temperature for 3–6 hours and overnight at 4 °C, then stored in PBS with 0.1% sodium azide. For imaging, brains were placed in 4.5% oxidized agarose (made by stirring 10 mM NaIO_4_ in agarose), transferred to a phosphate buffer solution, and placed in a grid-lined embedding mold for standardized orientation in an aligned coordinate space. Multiphoton image acquisition was accomplished by using the TissueCyte 1000 system (TissueVision, Cambridge, MA) coupled with a Mai Tai HP DeepSee laser (Spectra Physics, Santa Clara, CA), as described in Oh et al. (2014) for the Allen Mouse Brain Connectivity Atlas.

### Evaluation of tracer injection sites

Locations and extent of the injection sites in Sub and/or PS were evaluated based on the boundaries defined along DV axis of hippocampus in this study (see Figure 3). In addition, the term “effective injection site” was introduced in Cre-dependent viral tracing. Specifically, for example, when a tracer injection was involved in both region A and adjacent region B but the gene driving Cre was only expressed in cells of region A, then region A is the effective injection site because cells in region B would not express the GFP fluorescent tracer (see Table S2 for detailed Cre-lines and related effective injection sites).

### Projection quantification

Quantification of projection density was performed according to the methods and Informatics Data Processing Pipeline (IDP) for the Allen Mouse Brain Connectivity Atlas (Kuan et al., 2015; Oh et al., 2014;). Briefly, an alignment module of the IDP was used to align all injection experiments with the average 3D model brain after image reprocessing. A signal algorithm, based on a combination of adaptive edge/line detection and morphological processing, was applied to each section image to differentiate positive fluorescent signal from background signal. Segmented signal pixels were counted as projection strength in the claustrum and cortical areas. It should be noted that the detection algorithm operates on a per-image basis and that passing fibers and axon terminals were not distinguished (see e.g. Figure S10). Imperfect alignment of each injection image set with the Allen Mouse Brain CCF may also affect the quantification of the projections.

### Terminology used for mouse brain structures

The mouse brain atlas of Paxinos and Franklin (2012) and Allen Mouse Brain CCF (v3) were used in this study. Many structure terminologies are the same or similar in both atlases. See Table S3 for abbreviations/acronyms and full names of the brain structures used in this study.

## SUPPLEMENTAL INFORMATION

Supplemental information includes eleven figures and four tables.

## ACKNOWLEDGEMENTS

We are grateful for the technical support and expertise of the many staff members in the Allen Institute who are not part of the authorship of this paper. This work was funded by the Allen Institute for Brain Science. The authors wish to thank the Allen Institute founders, Paul G. Allen and Jody Allen, for their vision, encouragement, and support. The research was also supported by BRAIN Initiative Cell Census Network (BICCN) grant award U19MH114830 from the National Institute of Mental Health to H.Z. The content is solely the responsibility of the authors and does not necessarily represent the official views of National Institutes of Health and National Institute of Mental Health.

## AUTHOR CONTRIBUTIONS

Conceptualization: S.L.D. and H.Z. Transcriptomic data acquisition: K.A.S., T.N.N. and B.T. Connectivity data acquisition: K.E.H., P.B., K.N. and J.A.H. Data evaluation: Z.Y. and S.L.D. Clustering analysis: Z.Y., L.T.G. and O.F. Anatomical mapping of transcriptomic clusters: S.L.D. Connectivity data analysis: S.L.D. Figures preparation: S.L.D. Manuscript preparation: S.L.D., Z.Y. and H.Z., with inputs from other authors. Supervision: H.Z., B.T., J.A.H., E.S.L., J.W.P., C.K. and K.A.S. All authors read and commented on the manuscript.

## DELCLARATION of INTERESTS

The authors declare no competing interests.

---

**Table S1.** Transcriptomic clusters, gene makers and anatomic registration.

**Table S2.** Overall distribution of labeled axon terminals in the target regions of Sub and/or PS.

**Table S3.** Terminology and abbreviations used in this study.

**Table S4.** Comparison of the present and recent subicular studies.

**Figure S1.**
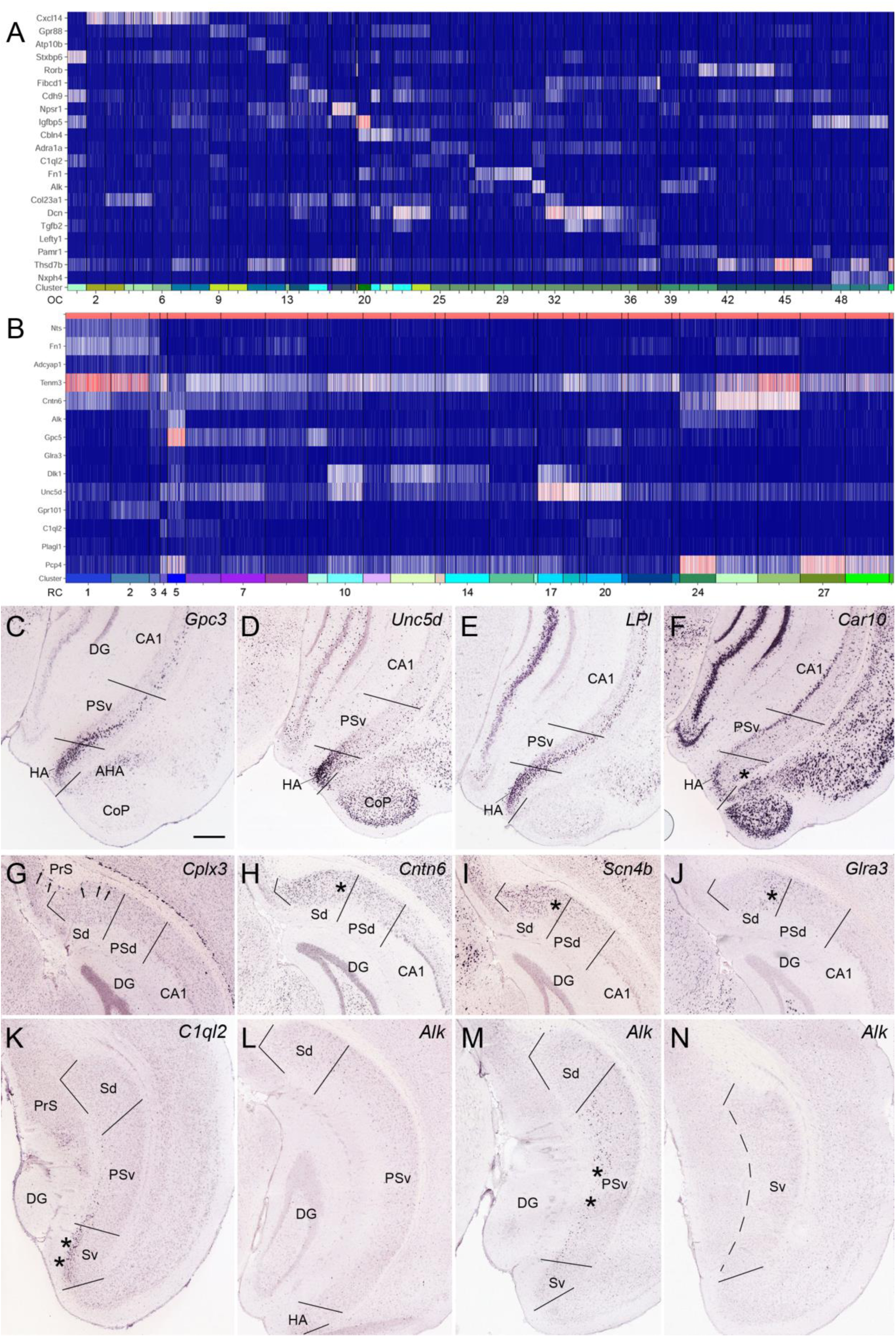
Heat map of selected gene markers from the original and re-clustering of transcriptomic data and ISH confirmation. Related to Figures 1 and 2. **(A)** Transcriptomic heat map of selected gene markers from original clustering (OC1-51). (**B)** Heat map of selected gene markers from re-clustering (RC1-29). (**C-N)** ISH confirmation of marker gene expression in sub-regions of HA (C-F), Sub (G-J) and the border region of Sub and PS (K-N). Strong and weak expression of *Gpc3*, *Unc5d* and *Lpl* was observed in HA and PSv, respectively (C-E). Note Car10 expression in superficial HA-py and HAL6 (* in F). In Spy, *Cntn6* (RC1) and *Scn4b* (RC2) are expressed in distal two-thirds while *Glra3* (RC3) in proximal one-third (* in H-J). Interestingly, *C1ql2* is expressed in the superficial Sv at the border with PSv (** in K; RC4) but not at further caudal levels. *Alk* is expressed in the superficial PSv at the border with Sv (** in M; RC5) but not at further rostral levels (L). Arrows in G point to layer 6b in Sub and PrS (OC49-50). Bar in C: 350µm (for C-N). For abbreviations see Table S3.

**Figure S2.**
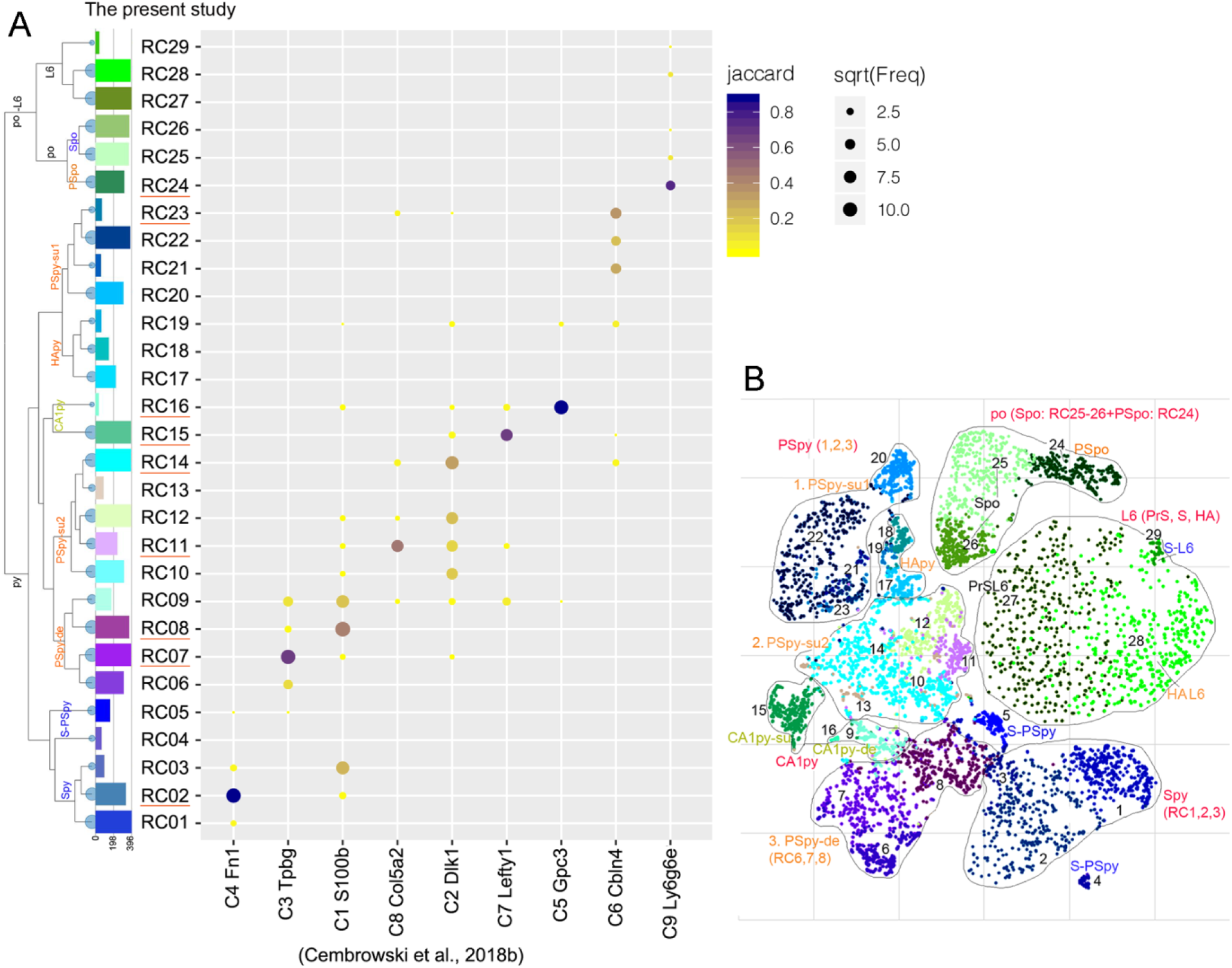
Confusion matrix and tSNE analysis. Related to Figure 1. (**A**) Comparison of the clusters identified in the present study and in Cembrowski et al. (2018b). This confusion matrix analysis indicates that the 9 clusters identified in Cembrowski et al. (2018b) partially map to the 29 clusters, mainly at subclass levels, identified in the preset study. See the Methods section for detailed workflow of this analysis. (**B**) Hierarchical and basic transcriptomic cell types shown in tSNE. Note that in PSpy, three major subclasses can be easily identified: PSpy-su1, PSpy-su2 and PSpy-de, and within these subclasses, multiple clusters are also found (for details of each cluster see Figure 1 and Table S1). For abbreviations see Table S3.

**Figure S3.**
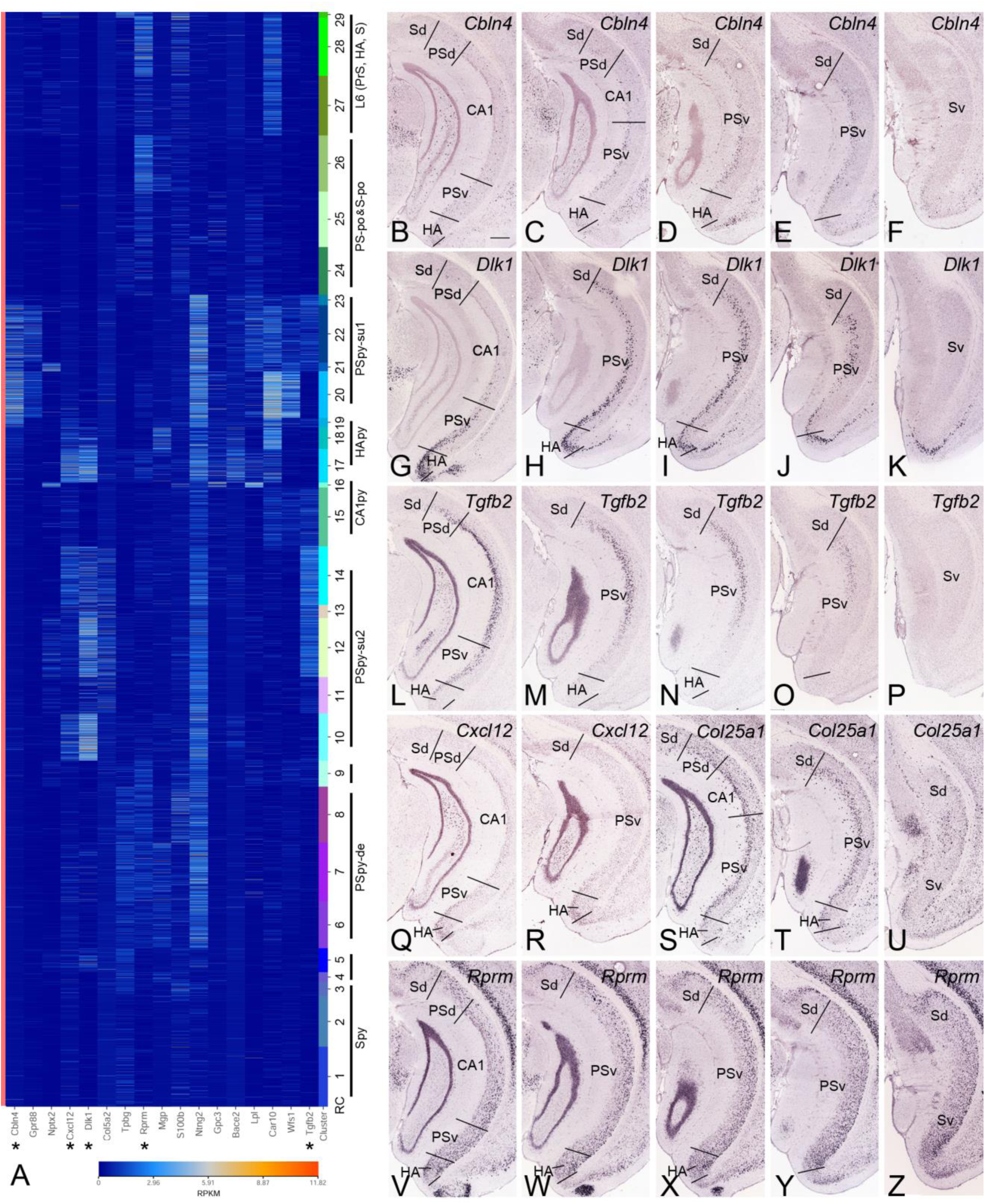
Marker genes for sub-types of PS and their spatial localizations. Related to Figures 1 and 2. **(A)** Heat map of selected marker genes expressed in subdomain of PS. Expression of selected genes (asterisks) in PS are shown in B-Z. (**B-Z)** ISH confirmation of marker gene expression in different sublayers of PS. (**B-F)** *Cbln4* expression in most superficial PSpy (see RC20-23 in A); (**G-K)** *Dlk1* expression in superficial PSpy (RC10-14) and HApy (RC17-18); (**L-P)** *Tgfb2* expression in most superficial and superficial PSpy (RC11-14 and RC21-23); (**Q-R)** *Cxcl12* expression in the most ventral part of superficial PSpy (RC14) and HApy (RC17-19); (**S-U)** *Col25a1*expression in superficial PSpy; (**V-Z)** *Rprm* expression in deep PSpy (RC6-8). Bar in B: 350µm (for B-Z). For abbreviations see Table S3.

**Figure S4.**
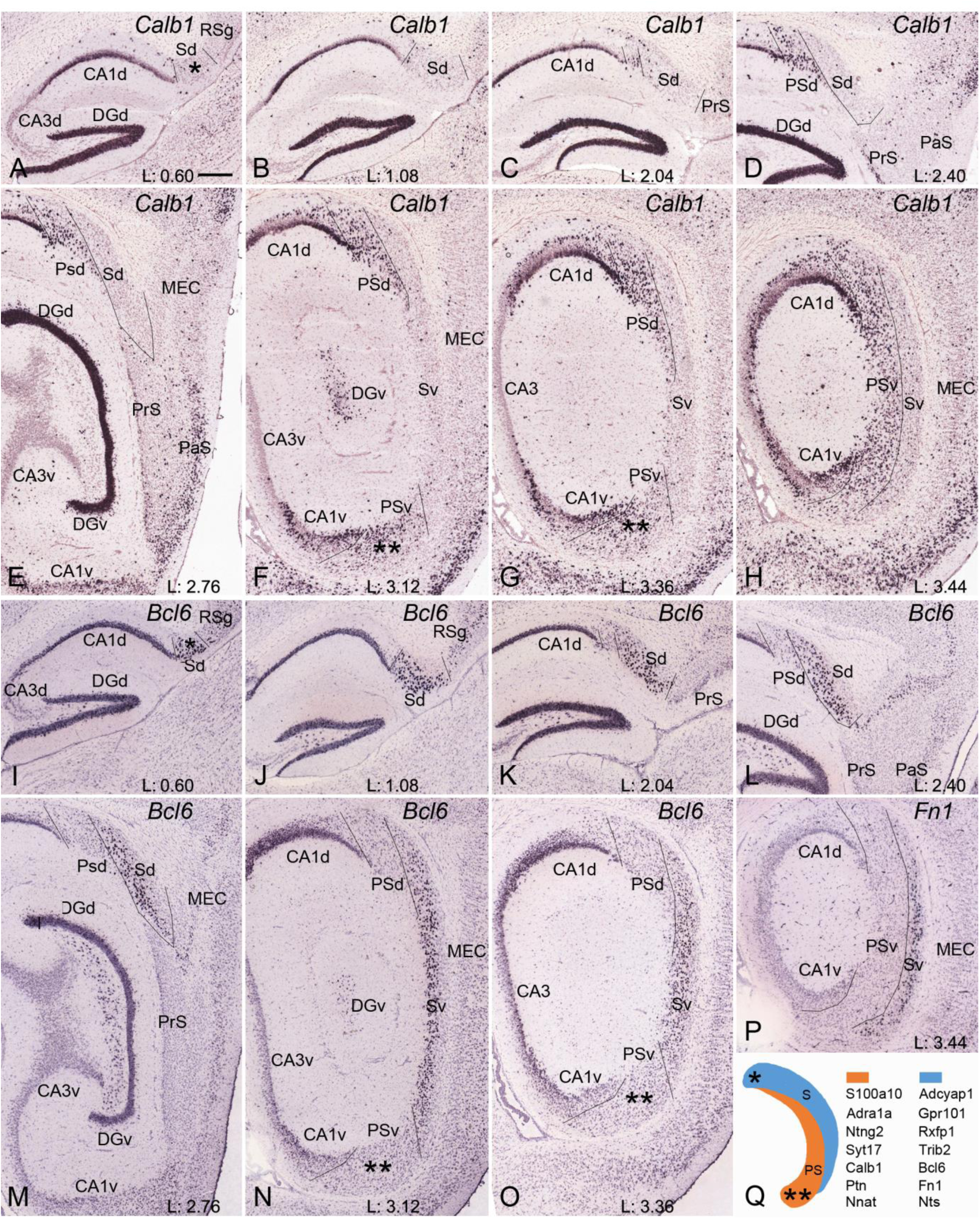
Borders, extent and topography of Sub and PS in sequential sagittal sections. Related to Figure 3. **(A-H)** Delineation of the borders of Sub, PS and CA1 from medial (A) to lateral (H) sequential sagittal sections stained for *Calb1* (PS marker). (**I-O)** Delineation of the borders of Sub, PS and CA1 from medial (I) to lateral (O) sequential sagittal sections stained for *Bcl6* (Sub marker). (**P**) *Fn1* ISH (another Sub marker) stain on a section corresponding to panel H. Note the complementary locations and different size (width) of the dorsal and ventral Sub and PS. (**Q)** A schematic drawing showing the DV difference and gradient of the size and extent of Sub (blue color) and PS (orange color) and representative genes selectively expressed in Sub and PS. Note that the most dorsomedial part (*) contains only Sub while the most ventrolateral part (**) contains only PS. Bar in A: 300µm (for A-P). Coordinates from Paxinos and Franklin (2012) are indicated at the bottom of each panel. For abbreviations see Table S3.

**Figure S5.**
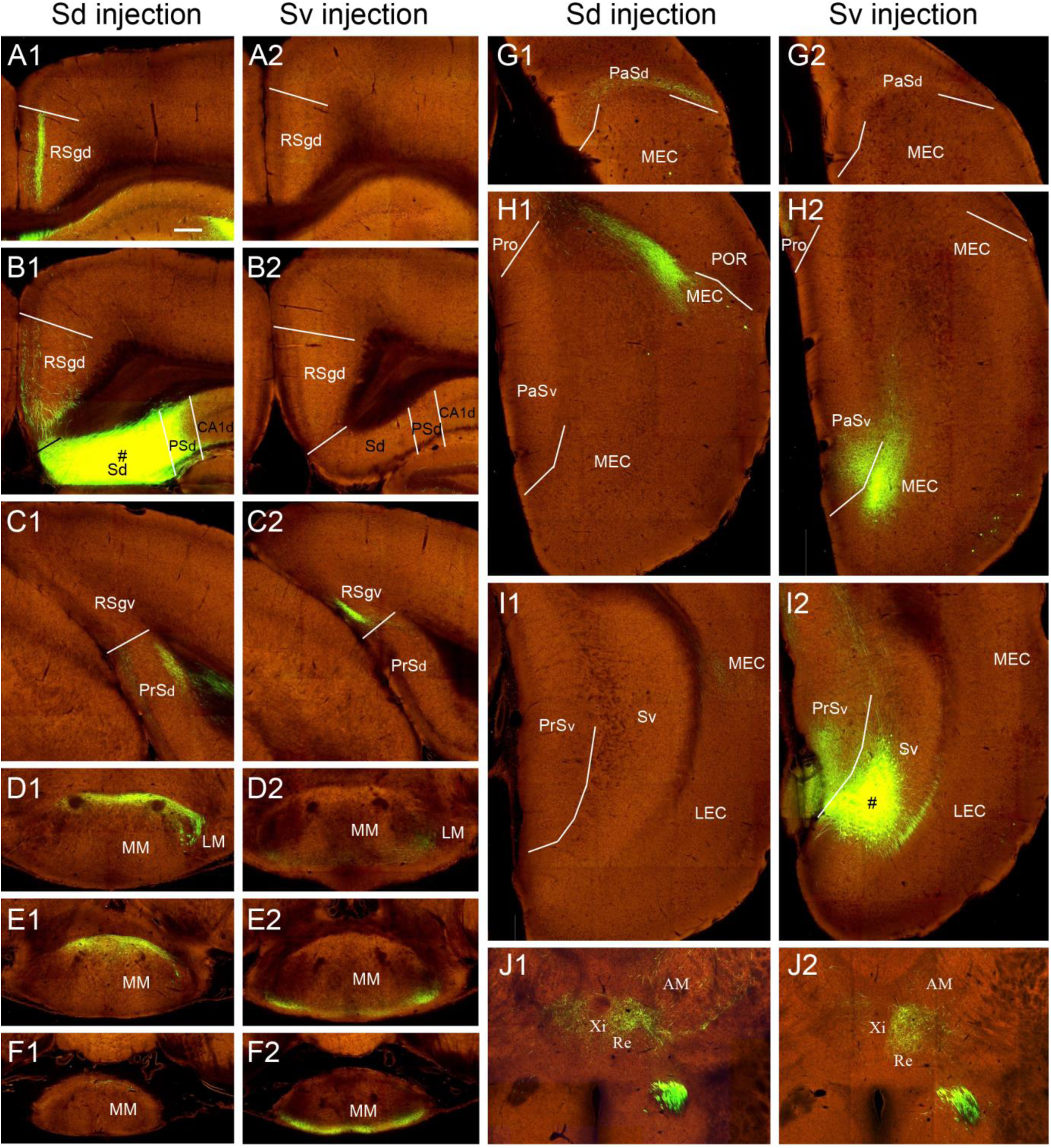
Comparison of the projections from Sd and Sv. Related to Figures 4 and 5. **(A1-I2)** Topographic projections from the most dorsal (A1, B1, …, I1) and the most ventral (A2, B2, …, I2) Sub. *Slc17a7*-IRES2-Cre (Figure 4A) for dorsal Sub injection and *Grik4*-Cre (another case ventral to Figure 4B) for the ventral Sub injection. The most dorsal Sub injection (in B1) results in terminal labeling in the most rostrodorsal RSg (A1), most dorsal PrS (C1), most dorsorostral MM (D1, E1, F1), most dorsal PaS (G1) and most dorsolateral MEC (H1, I1). In sharp contrast, the most ventral Sub injection (in I2) results in terminal labeling in the most caudoventral RSg (A2, B2, C2), most ventral PrS (I2), most ventrocaudal MM (D2, E2, F2), most ventral PaS (G2, H2), most ventromedial MEC (H2). A1 and A2, B1 and B2, …, I1 and I2 are sections at about the corresponding levels from the two Sub cases. (**J1 and J2)** Converging terminal labeling in the dorsal Re from both dorsal (J1) and ventral (J2) sub injections in two wild-type mice. Note no or few labeling is seen in the ventral Re, which instead receives projections from the PS (see Figure 6K). Bar: 233µm in A1 (for all panels). For abbreviations see Table S3.

**Figure S6.**
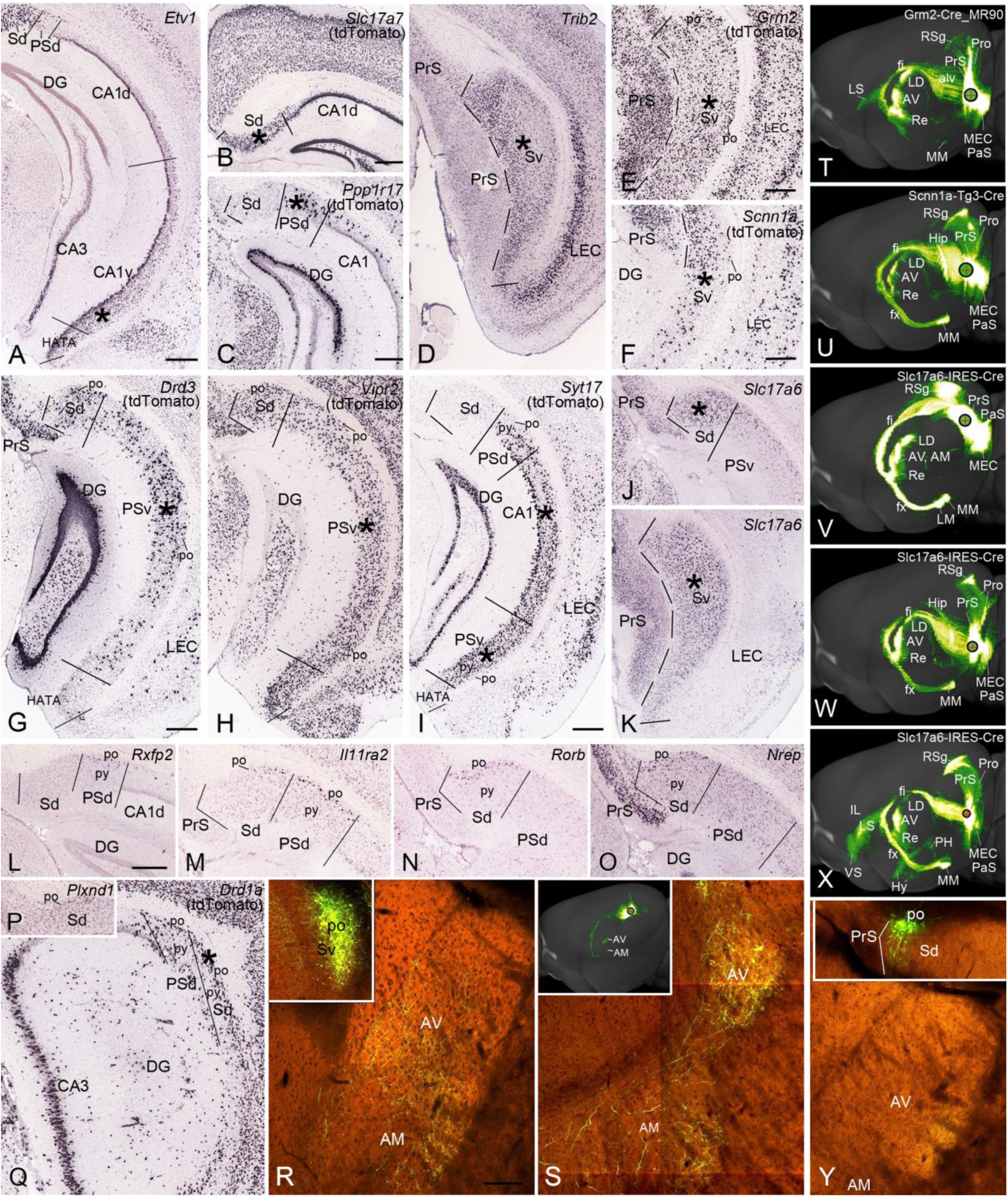
Cell type specific projections of Sub and PS. Related to Figures 4 and 6. Examples of mouse Cre-lines and related gene expression in injection sites of Sub, PS and CA1. Names of the genes (or genes whose promotors drive the Cre or tdTomato expression) are marked at top right corner of each panel. The asterisk in each panel indicates the center of the injection site. Overall projection patterns from these injections are shown in Figure 4 and Figure S6T-X. (**A-K)** Gene expression in related injection sites (indicated by asterisks). **(L-Q) G**ene expression in the polymorphic layer (po) of PS (L, *Rxfp2*; M, *Il11ra2*) and Sub (N, *Rorb*; O, *Nrep*; P, *Plxnd1*; Q, *Drd1a*). (**R and S)** Axon terminal labeling in AV and AM of the thalamus after rAAV injections in the po of Sv (Inset in R, *Plxnd1*-Cre_OG1 mouse) and Sd (Inset in S, *Drd1a*-Cre_EY262 mouse). (**T-X)** Overall projection patterns from additional 5 cases with injections involved in two or three regions: Sv and MEC injections in T, U and W; Sd and PrSd injection in V; Sv, PSv and MEC injections in X. Note that *Slc17a6*-IRES-Cre neurons in Sub do not appear to project to LS (V, W). In contrast, when the injection contains both Sub and PS strongly labeled terminals were observed in LS as well as VS and hypothalamus (X) with no or few labelling in IL and AON. (**Y)** An injection restricted in the most distal portion of Spo in Sd (Inset) of a *Slc17a7*-IRES2-Cre mouse resulted in axon terminal labeling only in the AV (*) with no labeling in AM. Interestingly, when an injection was placed in the most distal portion of Sd (both Spy and Spo) in a wild-type mouse, labeled axon terminals were only observed in AV, in addition to typical target labeling in RSg, MEC, PrS, PaS, Pro and MM (Table S2). The black circle with a red cross in each case indicates the injection site. Bars: 333µm in A and E; 395µm in B (for B, D, H), 420µm in C (for C and K); 300µm in F; 350µm in G (for G, I, J). 300µm in L (for L-Q); 140µm in R (for R, S and Y). For abbreviations see Table S3.

**Figure S7.**
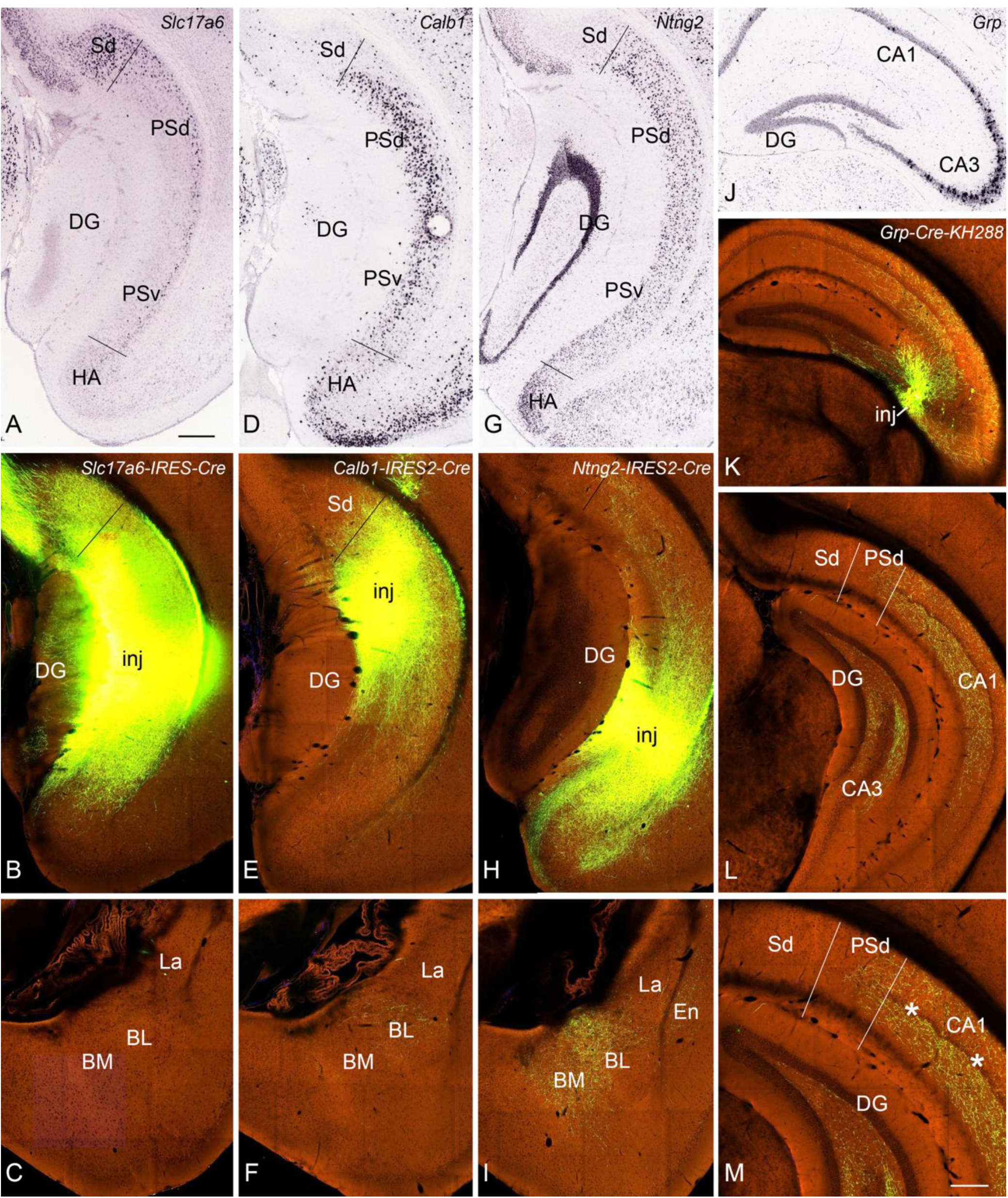
Cre-dependent anterograde viral tracing. Related to Figures 6 and 7. (**A-C**) A large injection in PS (“inj” in B) results in no terminal labeling in the amygdala (C) of a *Slc17a6*-IRES-Cre mouse, in which *Slc17a6* is expressed in Sub and deep PS (A). (**D-F**) An injection in PS (“inj” in E) results in terminal labeling in basolateral nucleus (BL) of the amygdala (F) of a *Calb1*-IRES2-Cre mouse, in which *Calb1* is expressed in the superficial two third of PS (D). (**G-I**) An injection in PS (“inj” in H) results in terminal labeling in both BL and BM (basomedial nucleus) of the amygdala (I) of a *Ntng2*-IRES2-Cre mouse, in which *Ntng2* is expressed in both superficial and deep PS (G). (**J-M**) A small injection in CA3 (“inj” in K) results in terminal labeling in both CA1 and PS of a *Grp*-Cre-KH288 mouse, in which *Grp* is expressed in CA3 (J). The image in M is a higher power view of the image in (L). The asterisks in (M) indicate the cell-dense band in the superficial pyramidal layer of CA1. Bars: 300 µm in A (for A-L); 200µm in M.

**Figure S8.**
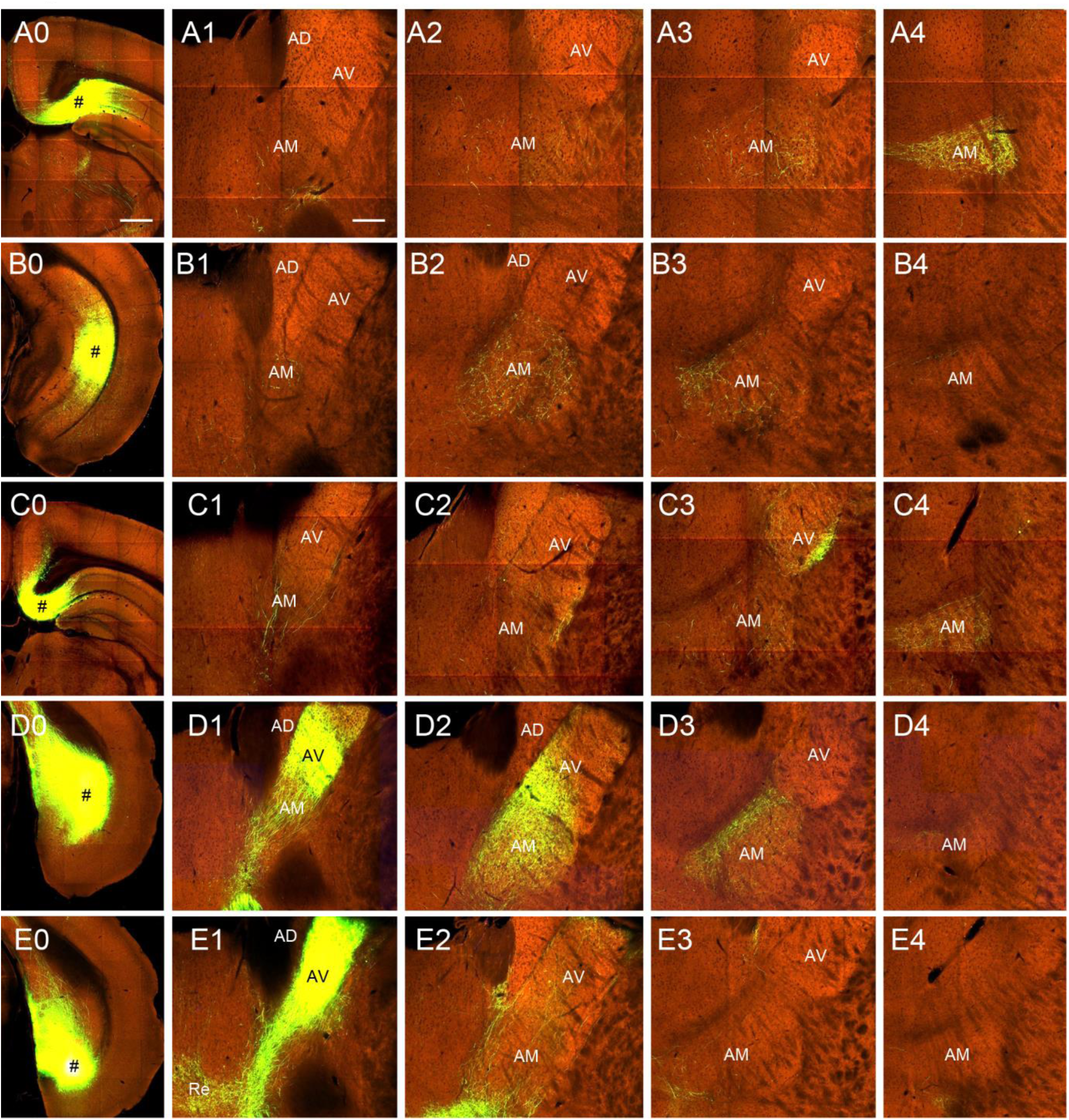
Topographic projections from Sub and PS to AV and AM. Related to Figures 4 and 7. (**A0-A4)** Projections from the most dorsal PS-CA1 (injection site # in A0) to AM (A1-A4) of a wild-type mouse. Labeled terminals are mostly seen in the most caudolateral AM (A3, A4). (**B0-B4)** Projections from PSv (injection site # in B0) to AM (B1-B4) in a *Drd3*-Cre_KI196 mouse. Labeled terminals are found in the rostromedial portion of AM (B2, B3). **(C0-C4)** Projections from the most dorsal Sub (injection site # in C0) to AV and AM (C1-C4) of a *Drd1a*-Cre_EY262 mouse. Labeled terminals are mostly seen in the most caudolateral AV and AM (C3, C4). (**D0-D4)** Projections from the ventral Sub (injection site # in D0) to AV and AM (D1-D4) in a *Trib2*-F2A-CreERT2 mouse. Labeled terminals are found in the rostromedial portion of AV and AM (D1, D2, D3). (**E0-E4)** Projections from the most ventral Sub (injection site # in E0) to AV and AM (E1-E4) in a wild-type mouse. Labeled terminals are found in the most rostromedial portion of AV and AM (E1, E2). Bars: 560µm in A0 (for A0-E0); 200µm in A1 (for all others). For abbreviations see Table S3.

**Figure S9.**
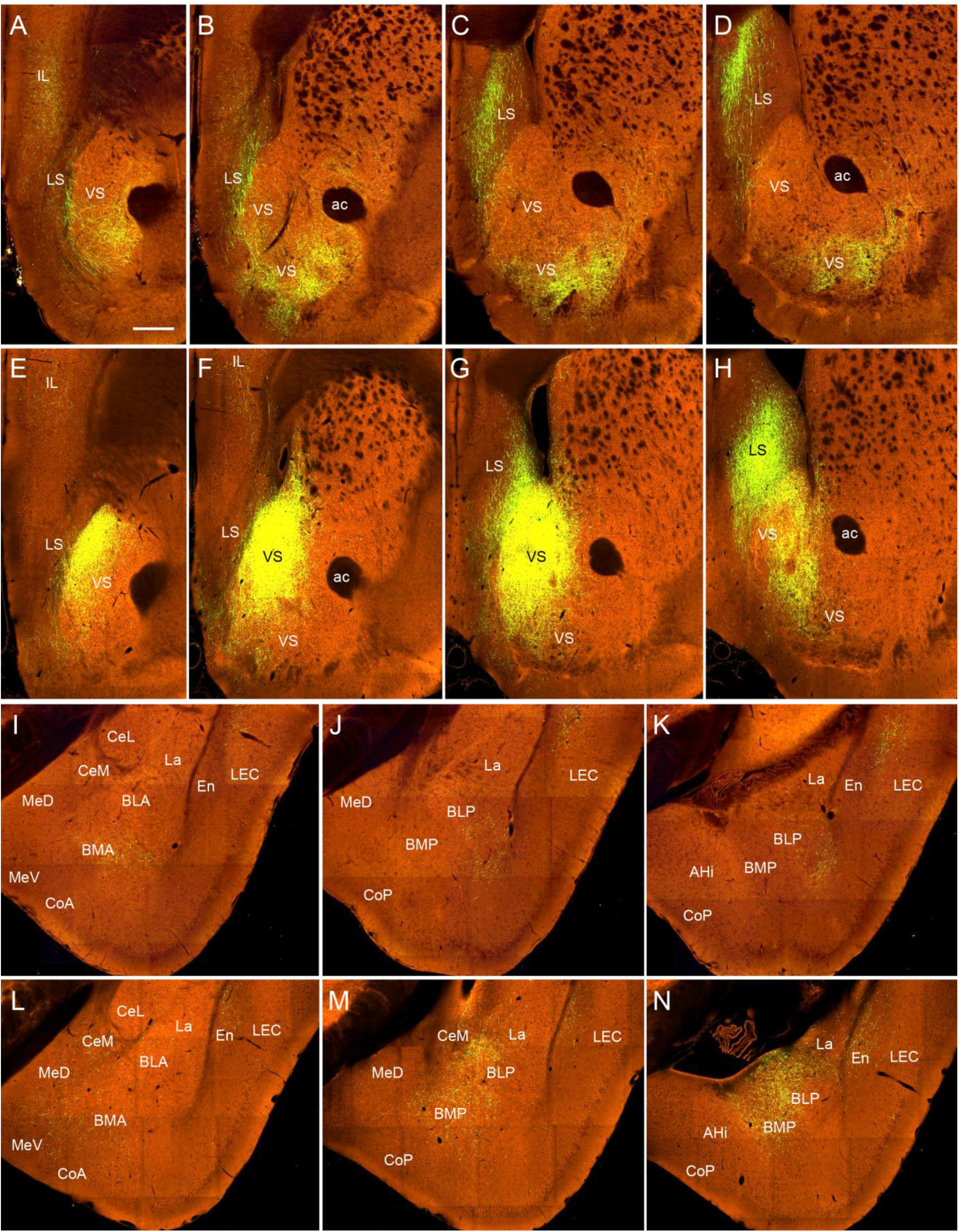
Topographic projections from PS to LS, VS and amygdala. Related to Figures 6 and 7. (**A-D)** Projections from PSd to LS and VS of a *Ppp1r17*-Cre_NL146 mouse. Labeled terminals are mostly seen in the dorsomedial LS (C, D) and lateroventral VS (A-D). (**E-H)** Projections from PSv to LS and VS of a *Ntng2*-IRES2-Cre mouse. Labeled terminals are mostly seen in the ventrolateral LS (G, H) and mediodorsal VS (E-H). (**I-K)** Projections from PSd to the amygdala and LEC of the *Ppp1r17*-Cre_NL146 mouse. Labeled terminals are clearly seen in the dorsal portion of LEC (I-K) as well as in the anterior portion of BM (BMA)(I) and the lateral portion of the posterior BL (BLP) (J, K) of the amygdala. (**L-N)** Projections from PSv to the amygdala and LEC of the *Ntng2*-IRES2-Cre mouse. Labeled terminals are clearly observed in the ventral portion of LEC (L-N) as well as in BMP and the medial portion of BLP (M, N) of the amygdala. La, Me and CeM also contain labeled terminals. Bar:350µm in A (for A-N). For abbreviations see Table S3.

**Figure S10.**
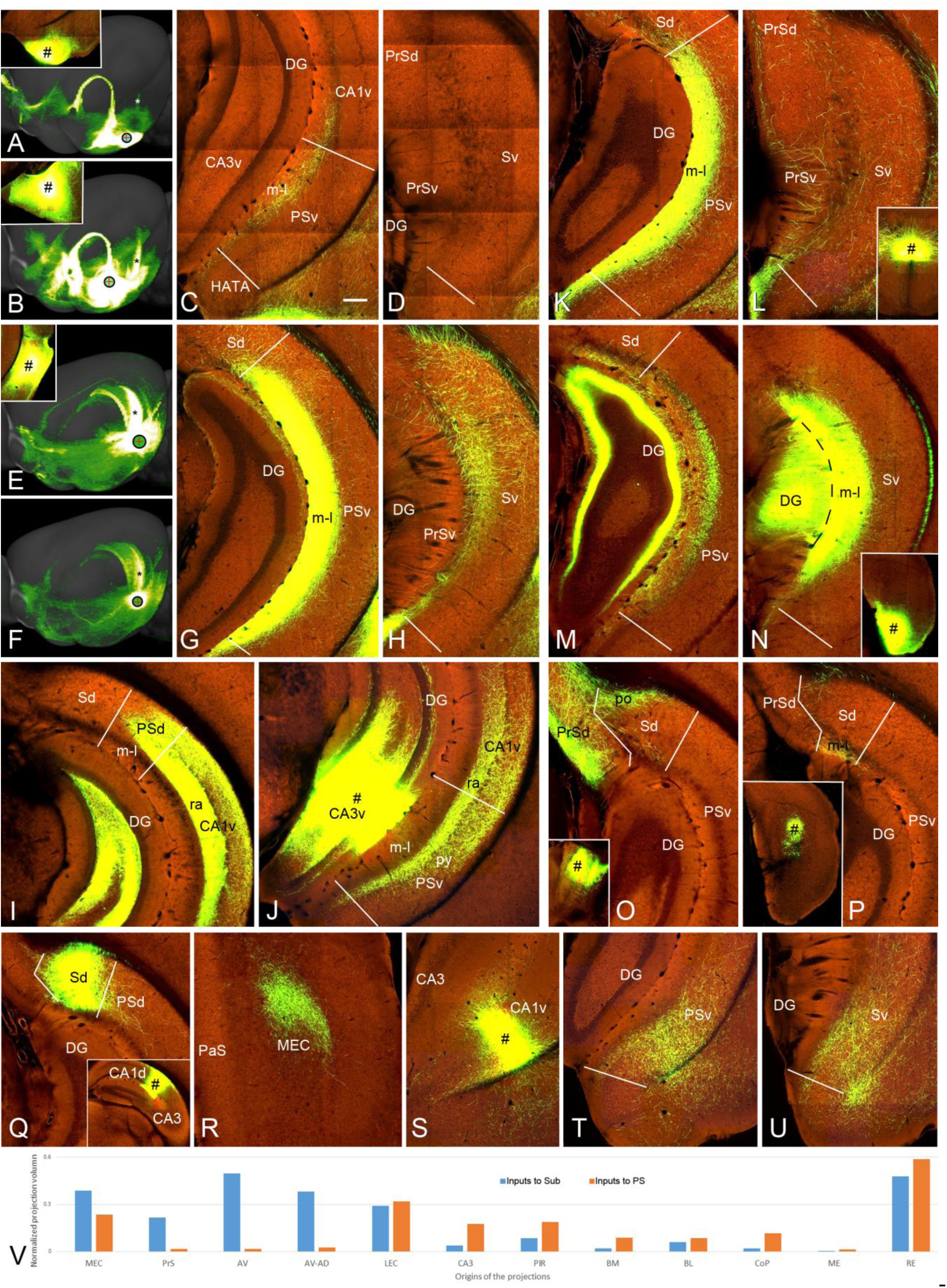
Comparison of the afferent projections to PS and Sub. Related to Figure 7. **(A-D)** Amygdaloid projections to PSv with no labeling to Sv. An injection in CoP (# in the inset of A) of a WT mouse results in clear terminal labeling in CA1v and PSv (see the asterisk in overall projection map in A and the labeled terminals in C) but not in Sv (D). An injection in BM (# in the inset of B) of a WT mouse results in strong terminal labeling in CA1v and PSv (see the asterisk in overall projection map in B). (**E-H)** LEC projections mainly target PS rather than Sub. An injection in LEC (# in the inset of E) of a *Cux2*-IRES-Cre mouse results in heavy terminal labeling in PSv (see the asterisk in overall projection map in E and the labeled terminals in G) but mostly fiber labeling in Sv (H). An injection in LEC of an *Otof*-Cre mouse similarly results in strong terminal labeling in PSv (see the asterisk in overall projection map in F). (**I)** An injection in dorsal CA3 of a *Dlg3*-Cre_KG118 mouse results in strong terminal labeling in PSd but not Sd. (**J)** An injection in ventral CA3 (# CA3v) of a *Syt17*-Cre_NO14 mouse results in heavy terminal labeling in PSv but not in Sv (not shown). (**K, L)** An injection in Re (# in the inset in L) of a WT mouse results in strong terminal labeling in PSv (K) but few in Sv (L). (**M, N)** An injection in MEC (# in the inset of N) of a *Cux2*-IRES-Cre mouse results in heavy terminal labeling in Sv (see the labeled terminals in N) but mostly fiber labeling in PSv (M). (**O)** An injection in AV (# in the inset of O) of a *Gpr26*-Cre_KO250 mouse results in terminal labeling in the polymorphic layer (po) of Sd and Sv (not shown) but not in PSd or PSv. (**P)** A small injection in the dorsal MEC (# in the inset of P) of a *Pcdh9*-Cre_NP276 mouse results in terminal labeling in the molecular layer (m-l) of Sd but not in PSv. (**Q, R)** Strong terminal labeling in Sd (all layers) and MEC (layers 5-6) after a dorsal CA1 injection (inset in Q) in a *Gpr26*-Cre_KO250 mouse (See Figure 4I). (**S-U)** An injection in ventral CA1 (S) and resulting terminal labeling in all layers of PSv (T) and Sv (U). (**V)** quantitative comparison of the afferent projections to Sub (blue bars) and PS (orange bars) from different origins. Note that the labeling of both passing axon fibers and axon terminals in panels H and M was included in the quantitative analysis. Bar: 200 µm in C (for All panels except A, B, E, F). For abbreviations see Table S3.

**Figure S11.**
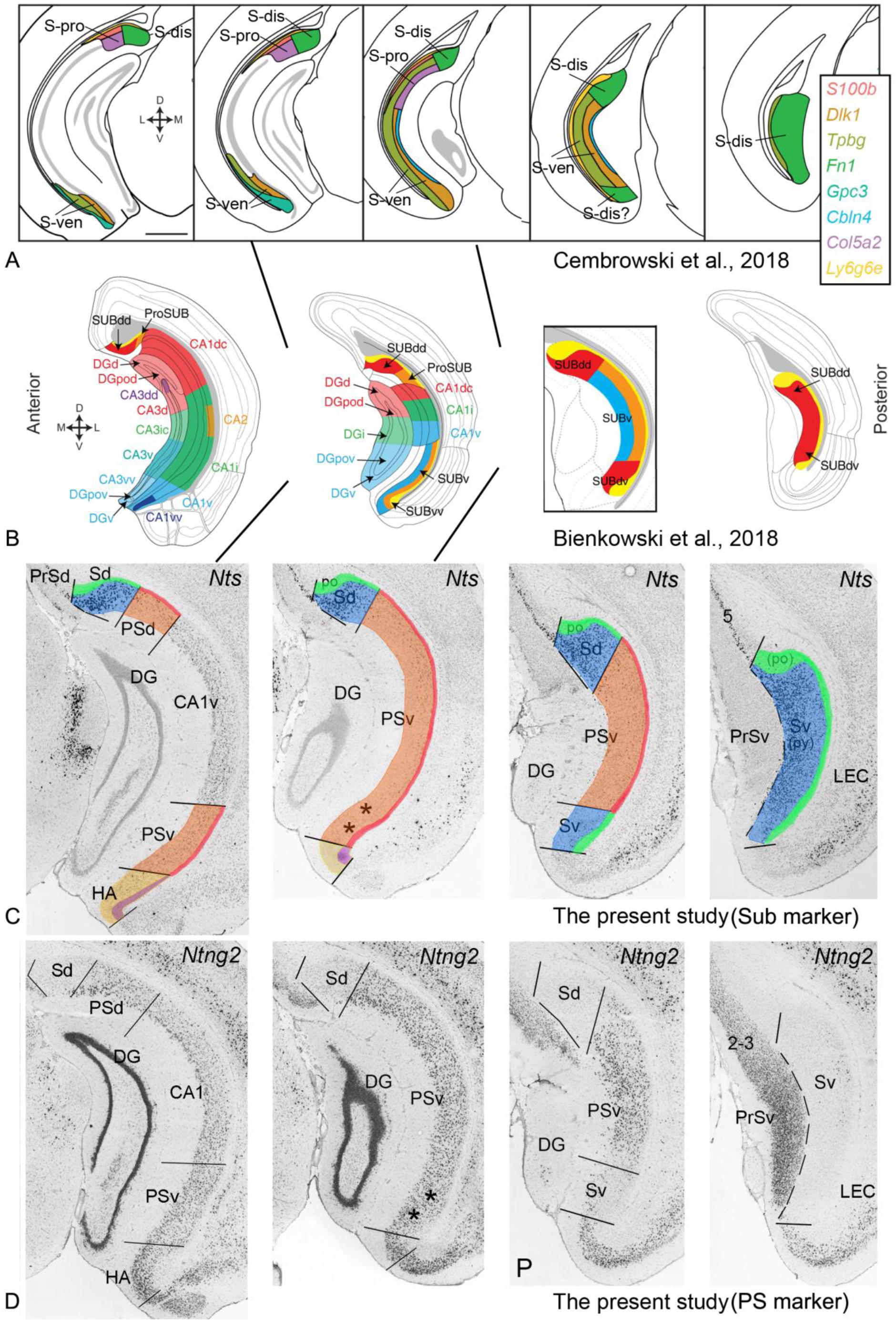
Summary and comparison of recently published studies (A, B) with the present study (C, D). Related to Figures 1, 3 and 7. It is clear that in the dorsal part, distal and proximal “Sub” (S-dis and S-pro in A), or SUBdd and ProSUB (in B), or Sd and PSd (in C and D) can be consistently identified across research groups. However, in the ventral part, major difference exists between the previous (A, B) and the present (C, D) studies. In A, the ventral “Sub” (S-ven) could not be subdivided into distal and proximal parts. In B, the ventral “Sub” could be subdivided into “SUBv” and “SUBvv”, in contrast to the dorsal part, which was subdivided into SUB and ProSUB (i.e., PS). In the present study, the ventral “Sub” is mostly occupied by PSv and HA (see the left two panels in C and D). The ventral Sub we define is located only at the caudal levels (see the right two panels in C and D). Our reason for this segmentation is based on the fact that the PSv region expresses marker genes of PSd (e.g. *Ntng2* and *S100a10*) but not those (e.g., *Nts* and *Bcl6*) of Sd (or S-dis in A or SUBdd in B). The PS and Sub re-defined in the present study are found to have differential connectivity, cell types and functional correlation, as demonstrated in detail here. In addition, since S-dis and S-pro were never clearly defined in previous studies it is impossible to simply replace PSv with ventral S-pro or replace Sv with ventral S-dis. Thus, the dorsal and ventral parts of Sub and PS we define here enable consistency at transcriptomic, connectional and functional levels along the entire DV axis.

